# Chronic Alcohol Drinking Drives Sex-Specific Differences in Affective Behavior and Medial Prefrontal Cortex Activity in CRF1:Cre:Tdtomato Transgenic Rats

**DOI:** 10.1101/2022.11.14.516444

**Authors:** SG Quadir, GM Arleth, MG Cone, MW High, MC Ramage, DP Effinger, M Echeveste-Sanchez, MA Herman

## Abstract

Alcohol use disorders (AUDs) are characterized by compulsive alcohol use, loss of control over intake, and a negative emotional state during abstinence. While AUDs are associated with both mood and chronic pain disorders, the relationship between these associations remains unclear. Corticotropin releasing factor-1 receptor (CRF1) has been implicated in alcohol (EtOH) use, affective states, and pain sensitivity; often in a sex-dependent manner. Using CRF1-cre transgenic rats, we found no sex differences in basal affective behavior with the exception of mechanical sensitivity, where females were more sensitive to mechanical stimuli. Following baseline testing, rats began EtOH (or water) drinking under intermittent access conditions. Females consumed more alcohol in the first week, but overall EtOH intake was not significantly different between males and females. Following 3-4 weeks of drinking, rats were tested again for negative affect. EtOH drinking decreased mechanical sensitivity, but no other group effects were observed. However, individual EtOH intake was directly correlated with anxiety- and depressive-like behavior in both sexes. Interestingly, EtOH intake inversely correlated with thermal sensitivity in males only. There were no group differences in CRF1+ neuronal activity in either prelimbic or infralimbic cortices, but final session EtOH intake was significantly correlated with activity in CRF1+ neurons in the infralimbic cortex. Together, our results suggest complex interplay between affective state, EtOH drinking, and the role of prefrontal cortex CRF1-containing neurons in mediating these behaviors. Additionally, these results highlight the importance of examining individual differences in AUD-related behaviors.

**SIGNIFICANCE STATEMENT:** Despite alcohol use disorders being extremely comorbid with mood and pain disorders, there is still a limited understanding of the interaction and directionality between the them. To investigate this problem, rats were tested for affective behavior before and after being allowed to drink alcohol for 6 weeks. While baseline behavior did not predict subsequent intake, alcohol intake predicted both anxiety- and depressive-like behavior. These findings were accompanied by increased activity of the corticotropin releasing factor 1 containing neurons in the infralimbic region of the prefrontal cortex. Together, these findings reveal a new mechanism for understanding alcohol use.

## 1. INTRODUCTION

In 2021, 28.6 million adult Americans met the criteria for having an alcohol use disorder (AUD)^1^. AUDs are generally characterized by excessive drinking and negative affective states upon drinking cessation^2^. These states can promote dependence as individuals drink to alleviate withdrawal symptoms such as increased anxiety and anhedonia^3, 4^ Conversely, anxiety also often precedes alcohol use and is linked to a more rapid progression of AUD in both humans^5–8^ and rodents^9–11^. AUDs and mood disorders are extremely co-morbid ^1, 12–14^; however, our understanding of this relationship remains incomplete as both are influenced by a variety of other factors including sex. For example, genetic models demonstrated that rats bred for high anxiety-phenotypes or congenital learned helplessness drink more alcohol (EtOH) than rats bred for low-anxiety or non-congenital learned helplessness phenotypes; but both differences are only present in females^15, 16^. Other studies found that anxiety-like behavior in the elevated plus maze predicts EtOH intake in male, but not female, rats^17^. In humans, AUD is more prevalent among men^18^; however, recent trends on EtOH consumption indicate that the prevalence of AUD among women is increasing at a faster rate than in men^19^ and the sex-related differences are narrowing^20^. Additionally, mood disorders are more common in women^21, 22^ and women are more likely to develop comorbid anxiety and AUD^23^. Together, these studies highlight a major gap in our understanding of AUDs: how sex influences affective behaviors that both precede and follow AUD development.

Affective behavior and AUDs are not only co-morbid in incidence, but also share similar neurobiological underpinnings. More specifically, both are mediated by the stress-related corticotropin releasing factor (CRF) system. While hypothalamic release of CRF results in activation of the hypothalamic-pituitary-adrenal axis, extrahypothalamic CRF acts as an important neuromodulator in the brain through activity at two G-protein coupled receptors, CRF1 and CRF2. In humans, polymorphisms in the gene encoding CRF1 are associated with AUD and AUD-related behaviors such as binge drinking, EtOH intake, and intoxication^24, 25^. Polymorphisms in CRF1 are also associated with affective behaviors such as post-traumatic stress disorder, panic disorder, depression, suicidality, and loneliness^26–32^. In rodents, systemically-administered CRF1 antagonists decrease anxiety-like behavior, EtOH intake, EtOH-induced negative affect and EtOH-induced hyperalgesia^33–39^. CRF1 is found extensively throughout the brain and has been studied in the context of both AUD and affective behaviors. However, most studies focus only on the classical stress regions, such the bed nucleus of the stria terminalis^40–42^ and the amygdala^41, 43–45^ despite CRF1 being abundantly expressed in cortical regions^46–48^. Indeed, alcohol preferring rats have increased CRF1 levels in the cingulate cortex, motor cortex, and somatosensory cortex^49^. CRF1 is also abundantly expressed in the medial prefrontal cortex,^50, 51^ but there are limited studies examining the effects of chronic EtOH drinking on prefrontal CRF1-containing neurons.

The present study uses behavioral testing and voluntary EtOH drinking in a CRF1 transgenic rat line to investigate the interaction between EtOH drinking and affective behavior in male and female rats. This study also examines the impact of chronic EtOH drinking on activity in the prefrontal cortex overall and specific to the CRF1 population.

## 2. METHODS

### 2.1 Subjects

32 adult male and female (n=16/sex) CRF1:cre: ^td^Tomato rats^52^ (8-11 weeks old, bred in-house) were housed in temperature and humidity-controlled rooms with 12h light/dark cycle (7:00 am lights ON). Rats had *ad libitum* access to food and water unless otherwise stated. Rats were handled before behavioral testing intake began. All animal procedures were performed in accordance with the [Author University] animal care committee’s regulations.

### 2.2 Splash Test

Rats were allowed to habituate in the behavioral testing room for 1h prior to testing. Rats were sprayed with 10% w/v sucrose solution, placed into a plexiglass/textured flooring behavioral testing chamber [50cm(*l*)×50cm(*w*)×38cm(*h*)] and recorded for 10min in red light conditions. Latency to groom and total time spent grooming were scored by an experimenter blind to experimental group using the Behavioral Observation Research Interactive Software (BORIS) program^53^. Latency to groom was used as a measure of anxiety-like behavior, whereas time spent grooming was used as a measure of depressive-like behavior. One subject was excluded from analysis due to equipment malfunction during testing.

### 2.3 Mechanical Sensitivity Testing (Von Frey)

Rats were brought into behavioral testing room 1h prior to testing and were habituated for 15min to a stainless-steel table [(78cm(*l*)×32cm(*w*)×19.5cm(*h*)] with a perforated sheet containing staggered holes (0.28cm diameter). Plastic filaments of increasing force (Bioseb EB2-VFF) were applied perpendicularly to hind paws until bending, and nocifensive responses defined as shaking, licking, or withdrawing the paw were recorded. Each paw was prodded 3 times/filament, beginning with the 2g filament and increasing until the rat exhibited a nocifensive response during at least 2 trials. Withdrawal thresholds for each hind paw were then averaged for each animal to determine the paw withdrawal threshold.

### 2.4 Novelty Suppressed Feeding (NSF)

Rats were given 1 Froot Loop in the homecage 48h prior to testing to prevent neophobia. 24h prior to testing food was removed from the cage. On test day, rats were habituated to behavioral testing room for 1h prior to testing. During testing, rats were individually placed into a brightly-lit (150 lux) behavioral testing chamber [50cm(*l*)×76cm(*w*)×40cm(*h*)] containing one Froot Loop on filter paper in the center. Latency to eat was recorded and used as a measure of anxiety-like behavior. After feeding initiation (or after 10min), rats were removed from the chamber and returned to the homecage where post-test consumption of pre-weighed Froot loops was recorded for 10min. Post-test consumption was used as a measure of motivated behavior.

### 2.5 Thermal Sensitivity Testing (Hargreaves Test)

Rats were assessed for thermal sensitivity using the Plantar Analgesia Meter (IITC). Rats were brought into the testing room 1h prior to testing and habituated to the heated glass (temperature: 32°C) apparatus for 20min. Infrared light (artificial intensity: 40) was focused onto each hindpaw and withdrawal latency was recorded. A maximum cutoff of 20s was used to prevent tissue damage. Each hindpaw was tested twice. If the latencies differed by more than 1s, then the paw was tested one additional time. Withdrawal latencies were averaged per paw then per animal.

### 2.6 Intermittent Access to Two Bottle Choice (IA2BC)

After 4d acclimatization to drinking out of two water bottles, one bottle containing 20% EtOH and one bottle of water were placed on cages for 24 hr on alternating days (MWF). Water drinking rats had their water bottles changed every 24h to ensure comparable treatment. Bottle placement was alternated to eliminate side preference. 20% v/v EtOH was prepared by diluting 95% EtOH (Pharmco Products Inc) in tap water.

### 2.7 Immunohistochemistry

24h after the last alcohol presentation, rats were anesthetized with isoflurane and transcardially perfused using phosphate-buffered saline (PBS) followed by 4% paraformaldehyde. Brains were post-fixed in 4% PFA at 4°C overnight and then transferred to 30% sucrose in PBS at 4°C until brains sank. Brains were serially sectioned at 40 μm and stored in cryoprotectant (30% sucrose, 30% ethylene glycol, 0.01% sodium azide in PBS) at 4°C.

For each animal, 4-6 sections containing the medial prefrontal cortex (ranging from 3.72 to 2.52mm anterior of bregma based on Paxinos & Watson^54^) were used. These 40 μm sections underwent PBS washes and were incubated in 50% methanol for 30 minutes. Slices were blocked in 3% hydrogen peroxide for 10min then 3% normal goat serum, 1% bovine serum albumin, 0.3% Triton X in PBS for 1h (with PBS washes in-between). Slices were incubated at 4°C for 48h with rabbit anti-cFos primary antibody (1:3000, Millipore Sigma ABE457) and mouse anti-RFP primary antibody (1:500, Invitrogen MA5-15257) in blocking solution. Slices were then washed with 0.1% Tween-20 in Trisbuffered saline (TNT) before 30min incubation in TNB blocking buffer (Perkin-Elmer FP1012). Next, slices were incubated with goat anti-rabbit horseradish peroxidase (1:400, Abcam ab6721) and goat anti-mouse 555 (1:800, Invitrogen A2123) in TNB buffer for 2h before TNT washes. Lastly, slices were incubated in tyramide conjugated fluorescein (Akoya Biosciences, NEL741001KT, 1:50) for 10 min. Following TNT washes, slices were mounted and sealed with mounting medium (Vector Laboratories H1500, Burlingame, CA). Slides were stored at 4°C until imaging with the Keyence BZ-X800 fluorescence microscope.

Imaging and quantification were performed by an experimenter blind to the condition and sex of each animal. Images were taken at 20X and stitched together using the BZ-X800 Analyzer program. One hemisphere was selected at random for quantification, where the area of interest was outlined and counted manually using the FIJI multi-point counter tool^55^. Cell counts were averaged per animal.

### 2.8 Statistical Analysis

All statistical analysis was performed using Graphpad Prism 8.0 (San Diego, CA). In all studies, the threshold for significance was set to *p*<0.05. Data was assessed for normality using the Shapiro–Wilk test or D’Agostino & Pearson. For data that did not follow a normal distribution, nonparametric tests were used as described below. Detailed statistics for each experiment can be found in **Table 1**, and are referenced by superscripts in the results section.

**Table 1.**
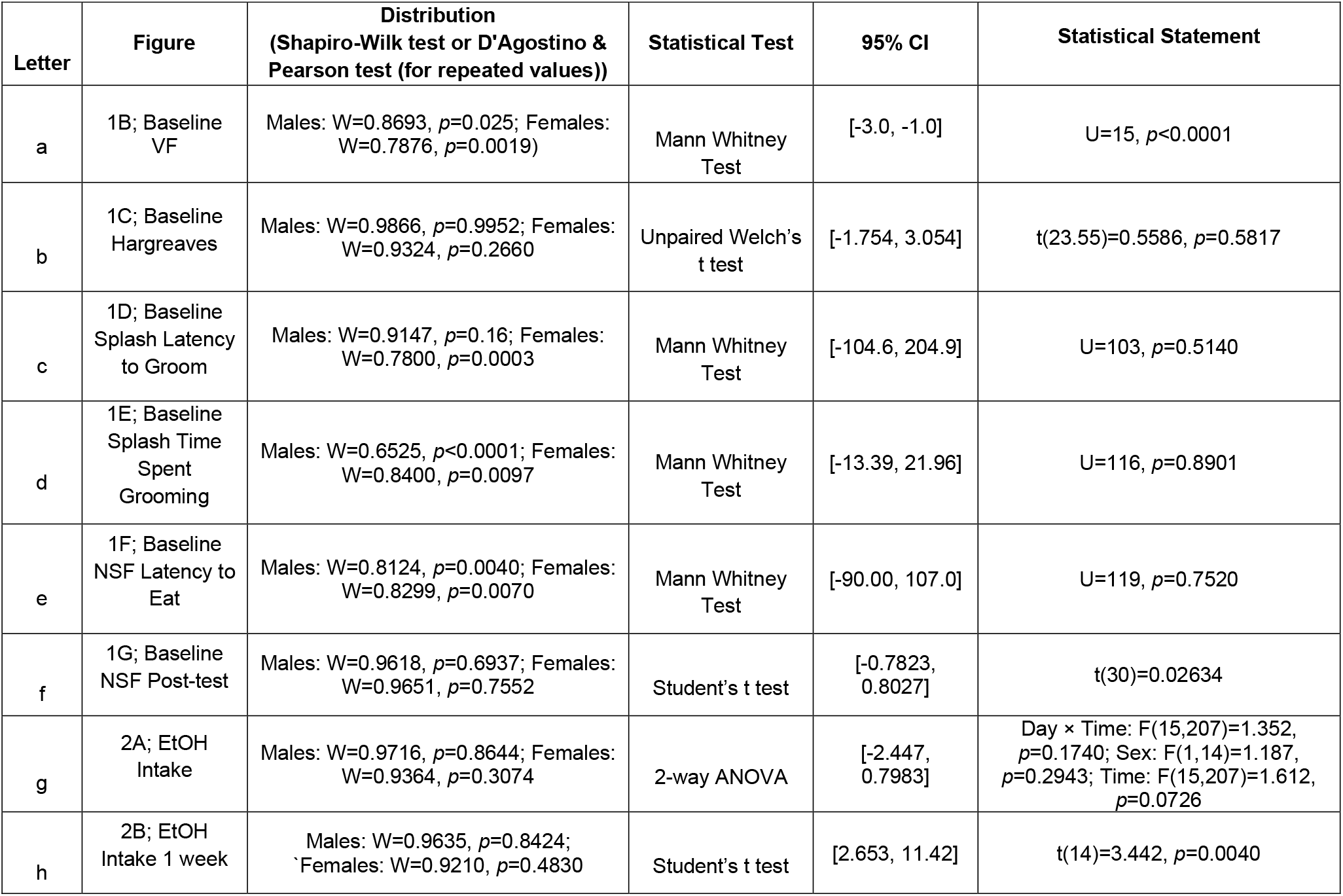
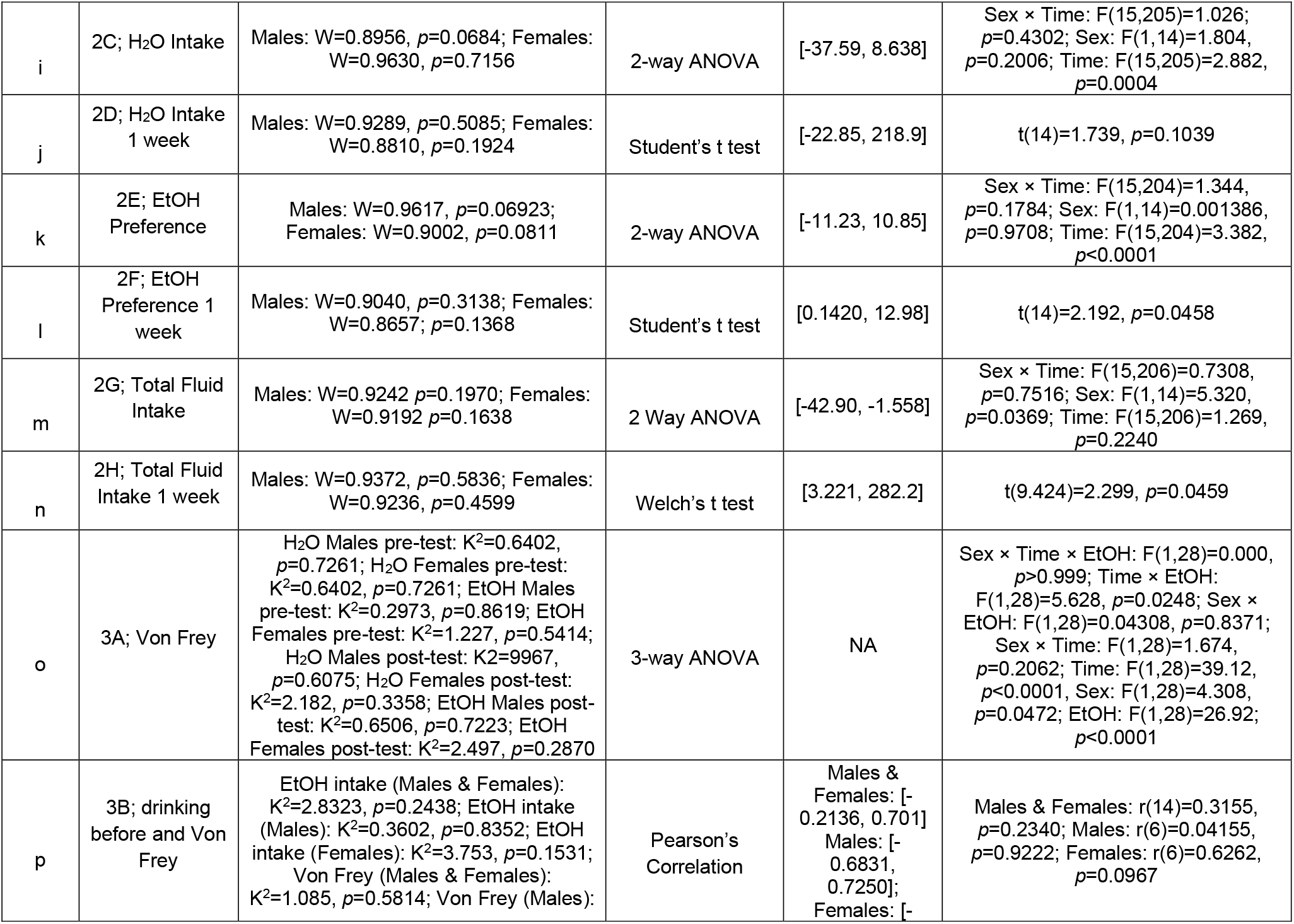
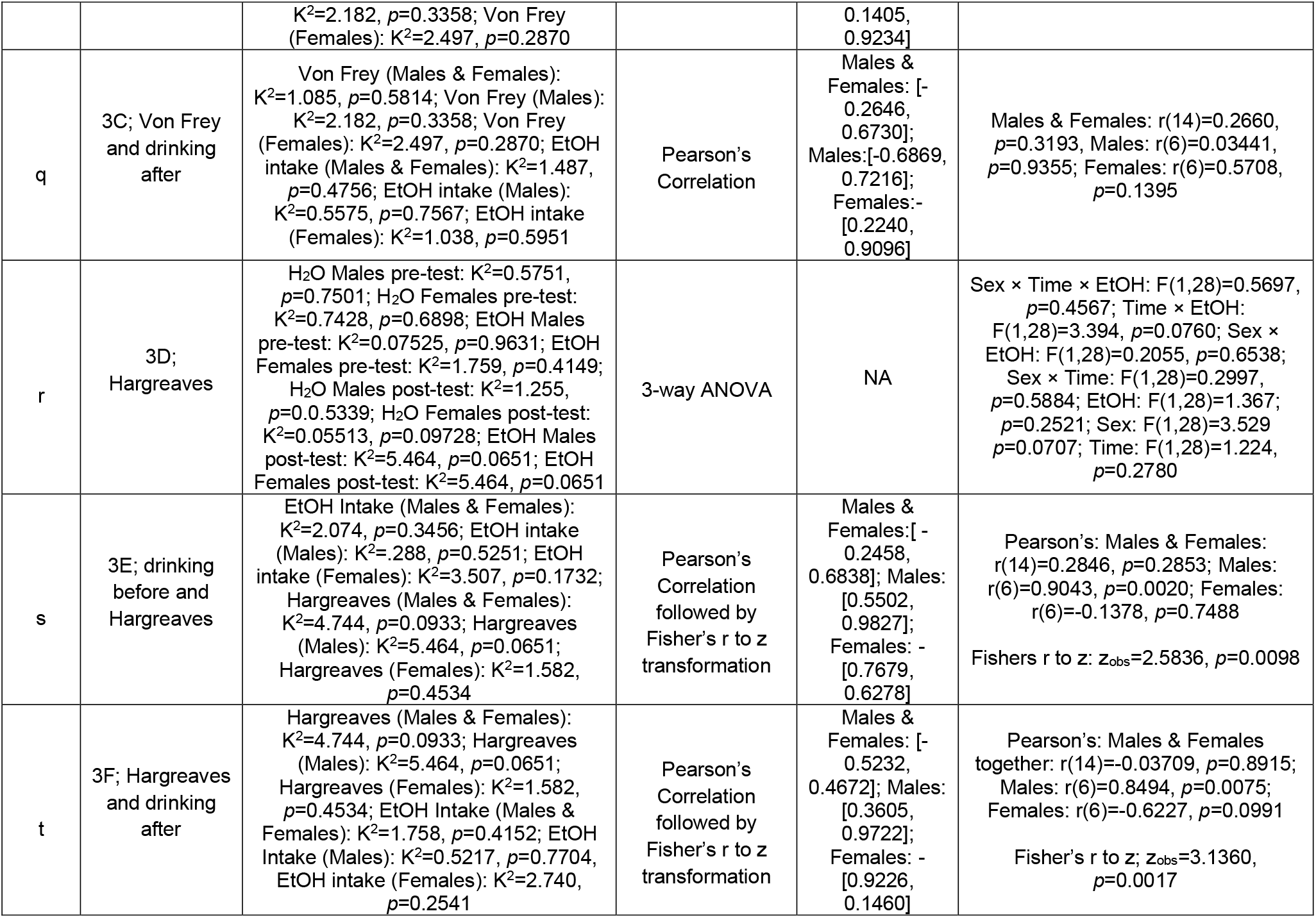
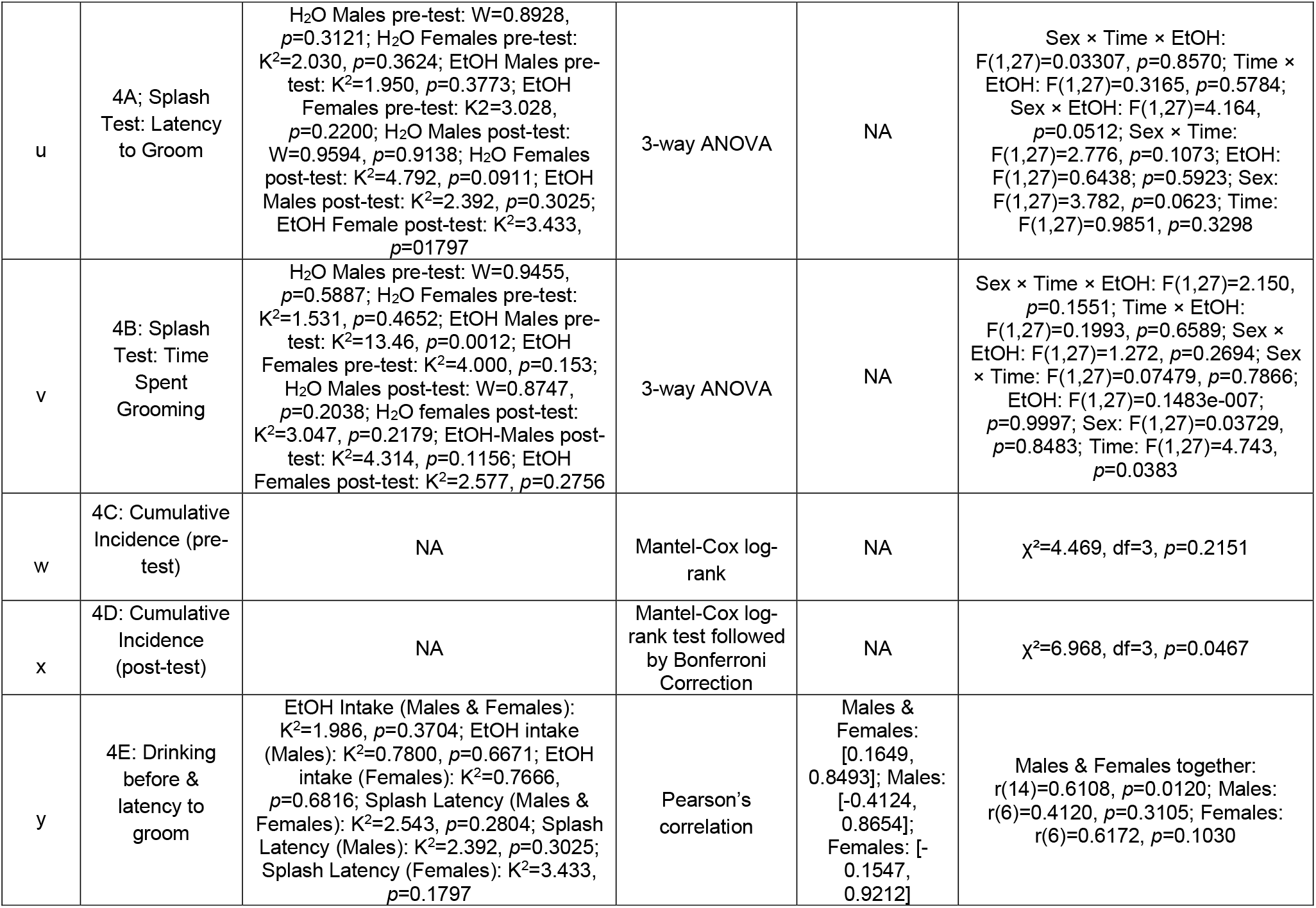
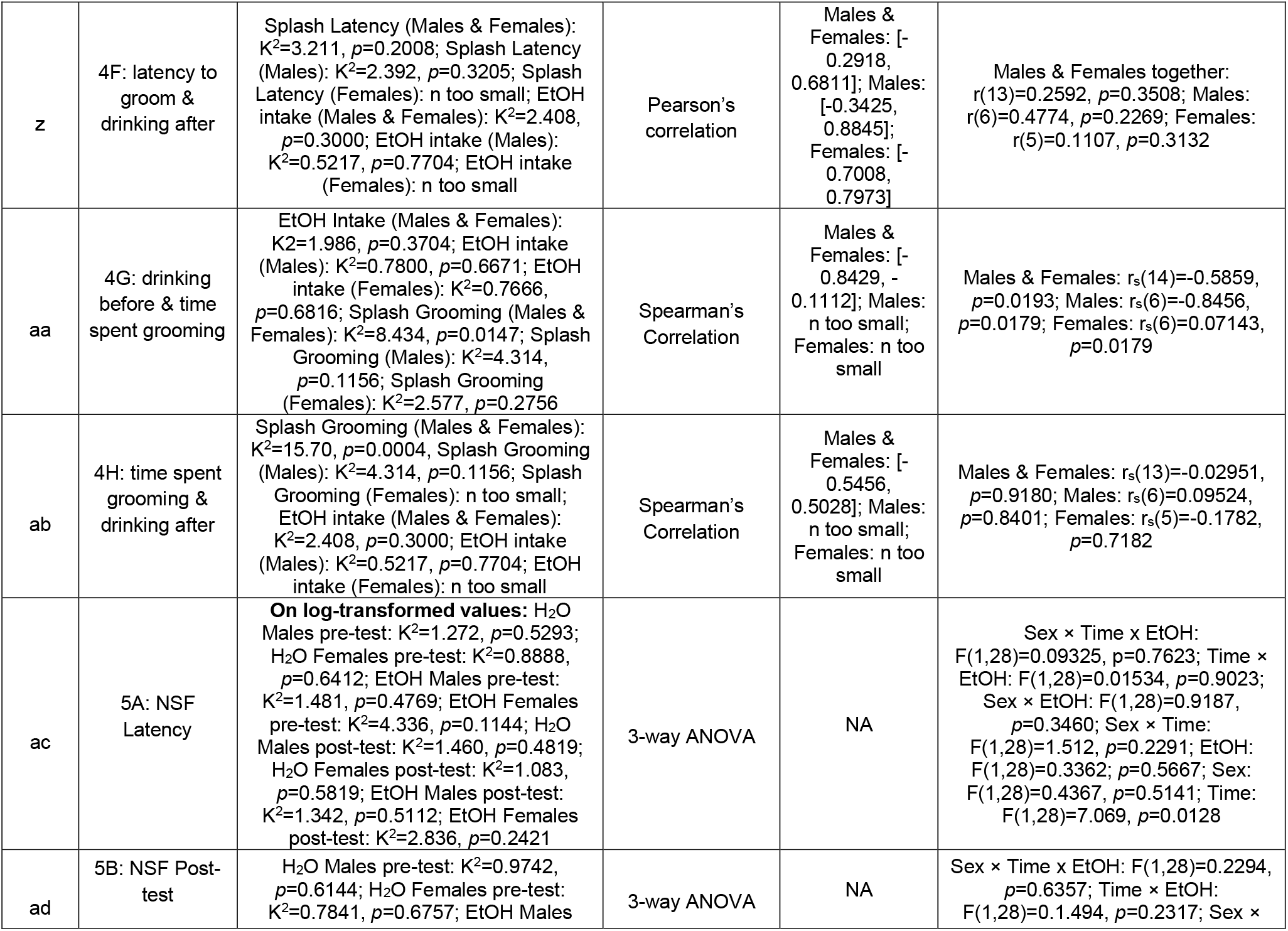
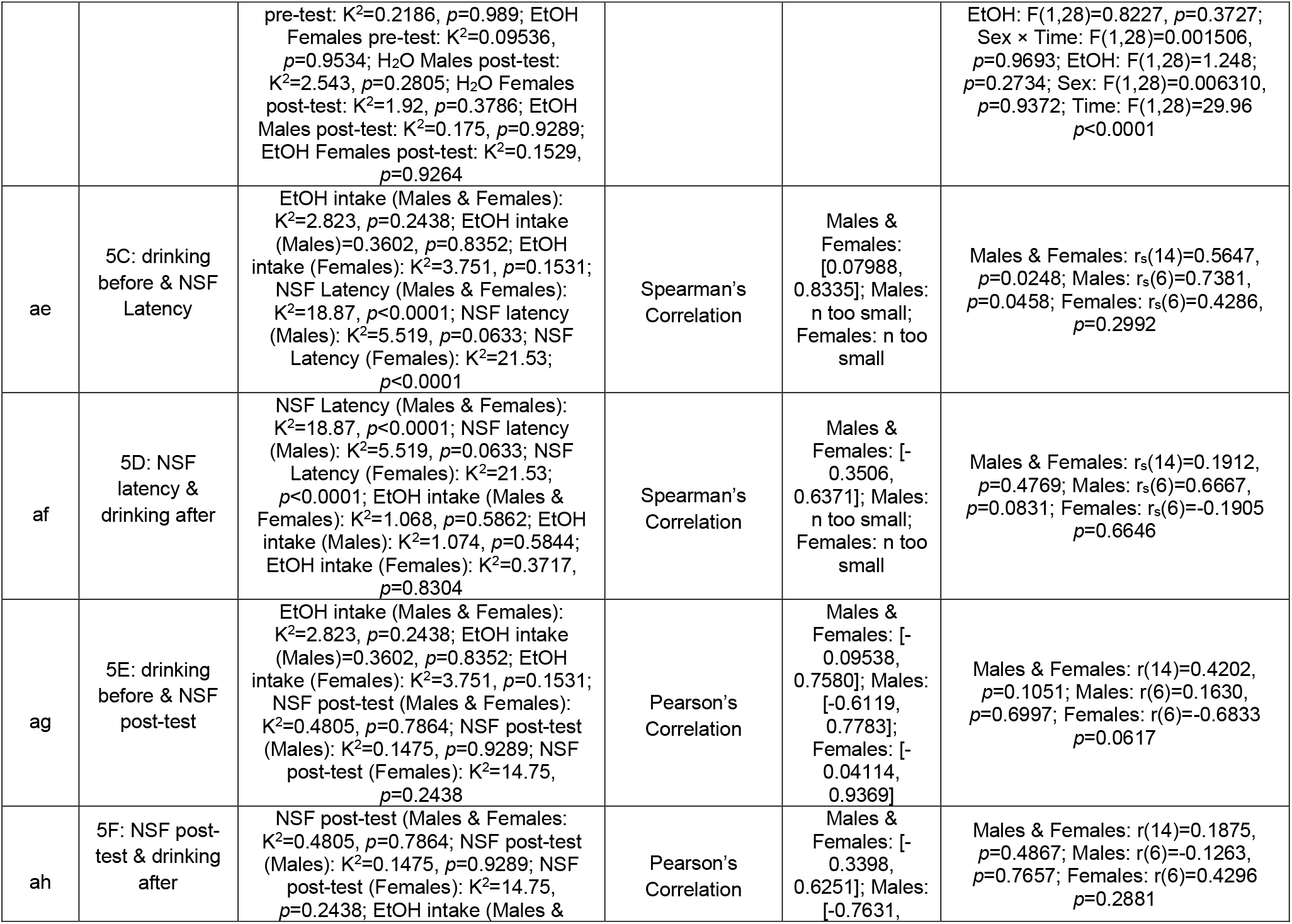
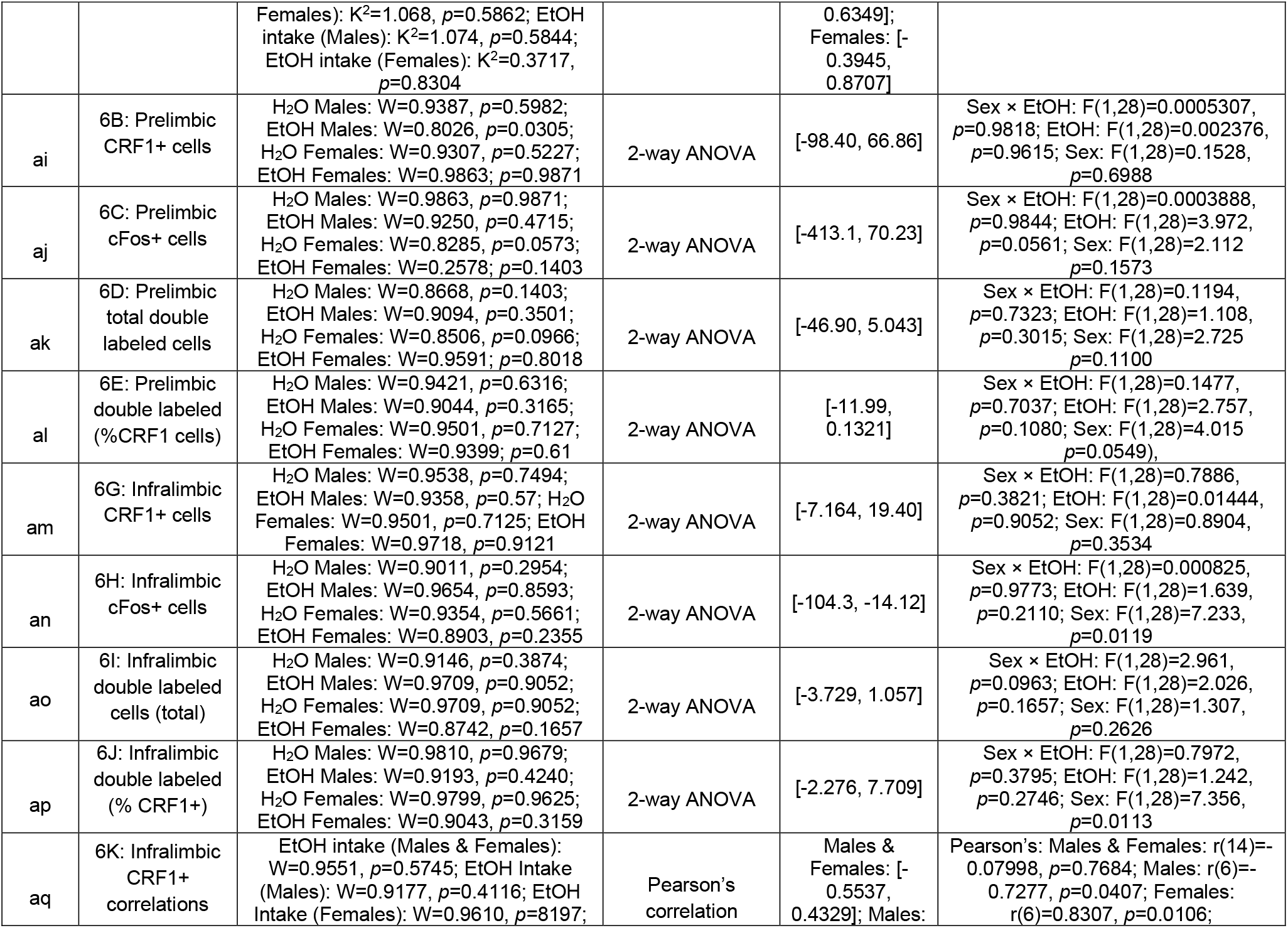
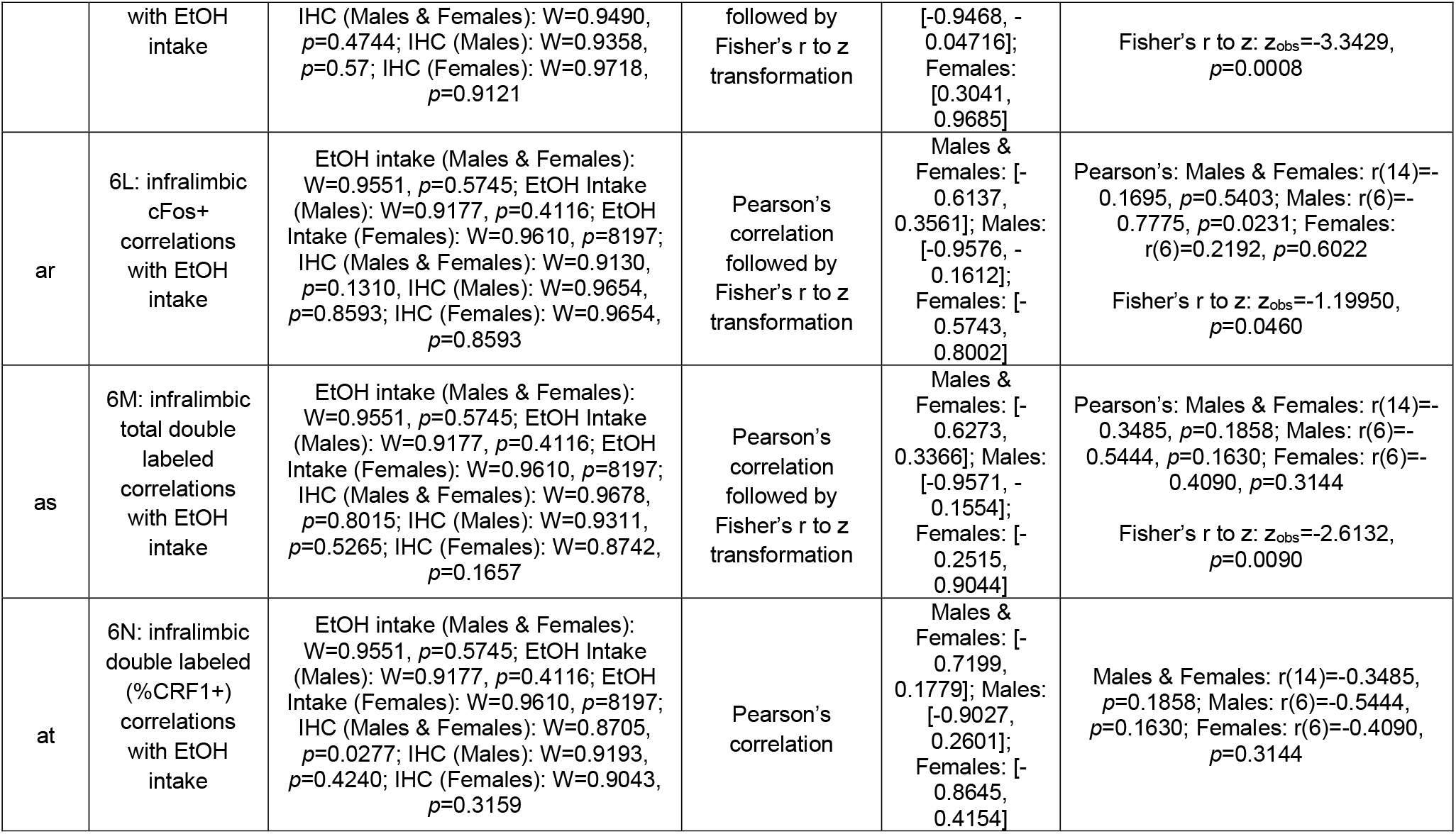
Statistics Table

Baseline behavioral tests were analyzed using an unpaired Student’s t test (parametric data with equal sample variances), Welch’s t test (parametric data with unequal sample variances), or a Mann-Whitney U test (nonparametric data). For cumulative incidence graphs, Kaplan-Meier survival curves were generated and were followed by Mantel-Cox log-rank test (followed by Bonferonni when applicable). Fluid intake and preference were analyzed by a mixed model 2-way ANOVA, using session as a within-subjects factor and sex as a between-subjects factor. Cumulative intake data were analyzed using an unpaired Student’s t test (equal sample variances) or an unpaired Welch’s t test (unequal sample variances). When behavioral tests were repeated (following drinking), data was analyzed using a 3-way mixed ANOVA (using time as a within-subjects factor while EtOH and sex were analyzed as between-subjects factors). When applicable, Bonferonni’s correction was used for multiple comparisons. Correlational analyses were performed using Pearson’s correlation (parametric data) or Spearman’s correlation (nonparametric data). To compare correlations between males and females, Fisher’s r to z transformation was used:

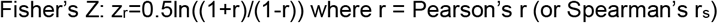

Z_r_ from males and females (denoted z_m_ and z_f_, respectively) was used to compute an observed z value, z_obs_ with the following calculation:

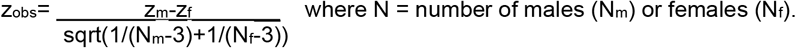

In each set of correlations, the same analysis (Pearson’s or Spearman’s) was used for both sexes to calculate a z_obs_.

Immunofluorescence data was analyzed by a 2-way ANOVA using sex and EtOH as between-subjects factors.

## 3. RESULTS

### 3.1 Experimental Timeline

As shown in **Fig. 1A**, rats were single-housed for 1 week prior to the onset of behavioral tests, which were conducted on separate but consecutive days. After baseline testing was complete, rats were left undisturbed for 3 days before being split into 2 groups, one allowed access to a bottle containing water and a bottle containing 20% EtOH under an intermittent access paradigm, and the other allowed access to only water bottles. After 3 weeks of drinking, post-drinking behavioral testing begun. These tests were separated by at least two drinking sessions in order to ensure testing did not disrupt EtOH consumption. During each test, rats were run in a pseudo-randomized subject order to minimize effects of circadian cycle and withdrawal time. After the last behavioral test, rats underwent a final EtOH drinking session. 24h after the final drinking session, rats were perfused and tissue was collected as described above.

**Figure 1.**
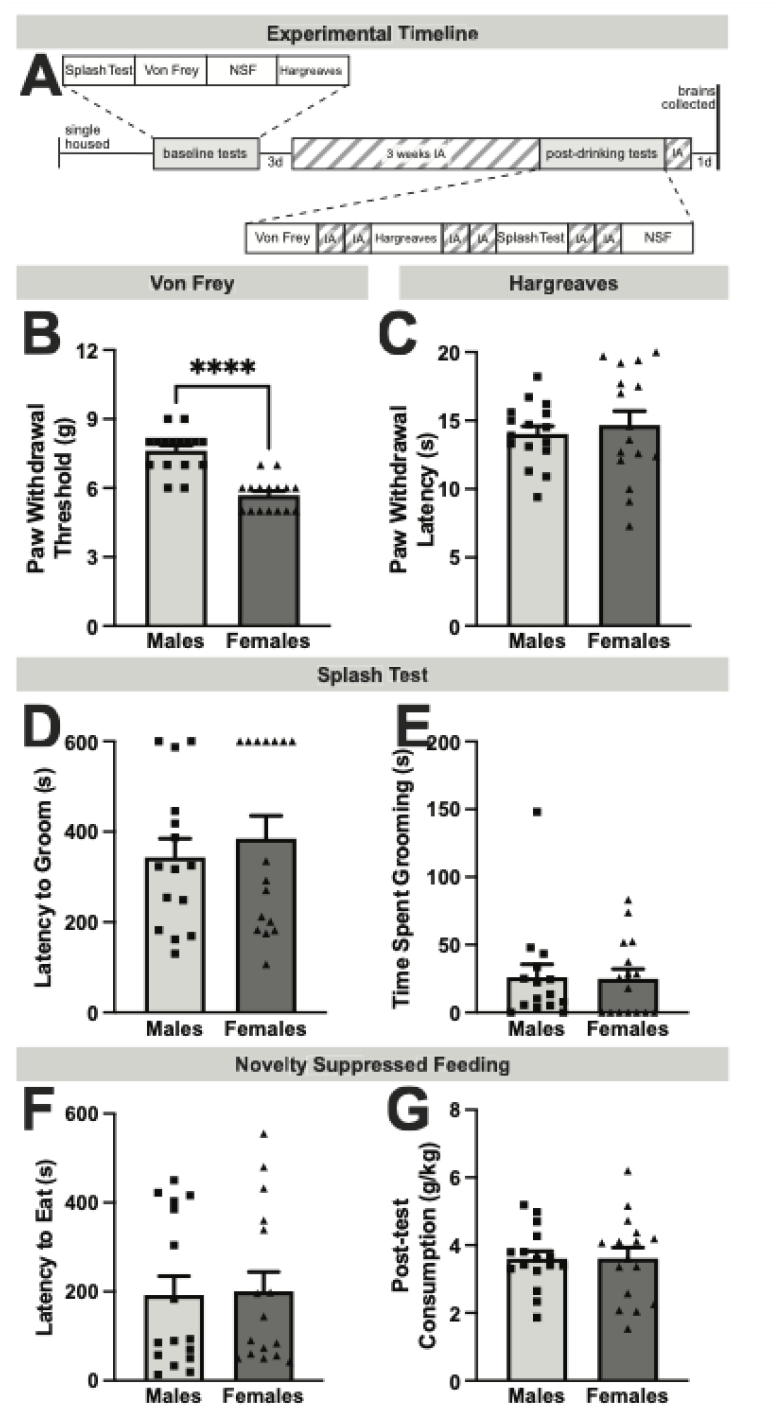
Experimental timeline and baseline behavior. **A**) Timeline of entire experiment. **B**) Mechanical sensitivity in Von Frey test, as measured by paw withdrawal threshold. **C**) Thermal sensitivity in the Hargreaves test, as measured by latency to withdraw. **D)** Anxietylike behavior in the splash test, as measured by latency to groom. **E)** Depressive-like behavior in the splash test, as measured by time spent grooming. **F)** Anxiety-like behavior in the Novelty Suppressed Feeding test, as measured by latency to eat. G) Motivation to eat in the Novelty Suppressed Feeding test, as measured by post-test consumption

### 3.2 Baseline Affective Behavior

Rats underwent a series of baseline behavioral tests to probe potential sex differences in behavior prior to EtOH drinking. In particular, we found sex differences in the Von Frey test of mechanosensitivity where female rats exhibited decreased withdrawal thresholds^a^, suggesting increased mechanical sensitivity (**Fig. 1B**). However, these differences were unique to mechanosensation, as there were no sex differences in the Hargreaves test of thermosensitivity^b^ (**Fig. 1C)**. Additionally, there were no sex differences in the Splash test parameters including latency to groom^c^ **(Fig. 1D)** or time spent grooming **(Fig. 1E)**^d^. Some animals never initiated grooming and were thus assigned artificial latency values of 600s; however, cumulative incidence of grooming initiation curves **(Extended Data 1-1)** revealed no differences between sexes. There were also no sex differences in latency to eat^e^ (**Fig. 1F**) or post-test home-cage chow consumption^f^ **(Fig. 1G)** in the NSF test. Similarly, there were no differences in cumulative initiation of feeding **(Extended Data 1-2)**. Together, these results indicate that while female CRF1-cre rats display increased basal mechanosensitvity, there are no sex differences in basal anxiety-like or depressive-like behavior.

**Extended Figure 1-1.**
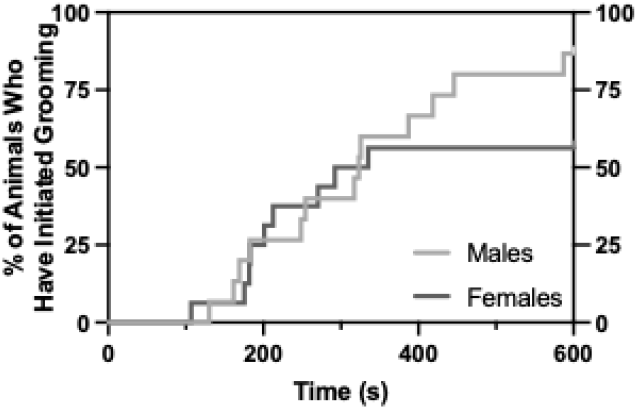
Cumulative Incidence of Grooming Initiation in the Splash Test. Even when accounting for the rats that never groomed (rather than artificially assigning them a maximum value or excluding them from studies), there was no sex difference in grooming initiation (Mantel-Cox log-rank test: χ^2^=1.430, df=1, *p*=0.2318). There was also no sex difference in total time spent grooming (**C;** Mann-Whitney test, *p*=0.8901).

**Extended Figure 1-2.**
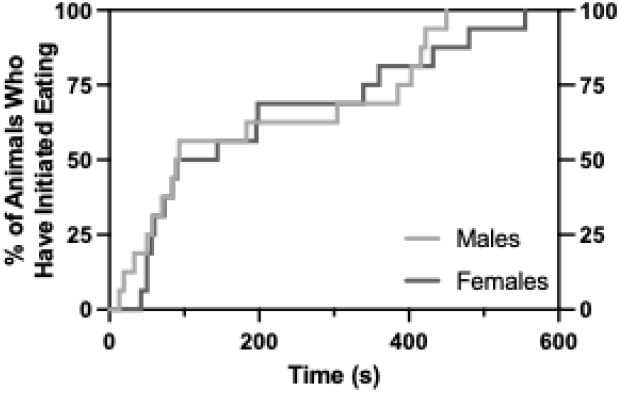
Cumulative Incidence of Feeding Initiation in the Novelty Suppressed Feeding Test. There was no effect of sex on cumulative occurrence of feeding initiation (**B**; Mantel-Cox log-rank test, χ^2^=0.3366, df=1, *p*=0.5618).

### 3.3 Voluntary EtOH Drinking

After assessing baseline affective behavior, rats were divided into two groups. The rats in the EtOH group were presented with 20% EtOH and water under an intermittent access two-bottle-choice procedure, whereas the rats in the water group received two water bottles. While there were no significant sex differences in overall EtOH intake^g^ (**Fig. 2A**); females consumed significantly more EtOH in the first week of drinking^h^ (**Fig. 2B**). Interestingly, there were no sex differences in water intake overall_i_ (**Fig. 2C**) or during the first week^j^ (**Fig 2D**), but water intake decreased over time in both sexes. We also looked at EtOH preference, calculated as volume of 20% EtOH consumed divided by volume of total fluid consumed during the session x 100%. While EtOH preference increased over time in a non-sex-dependent manner^k^ (**Fig. 2E**), females exhibited a stronger EtOH preference during the first week^l^ (**Fig. 2F**). Additionally, females consumed significantly more total fluid compared to males overall^m^ (**Fig 2G**) and during the first week^n^ (**Fig. 2H**). There were no significant correlations between any of the affective behaviors tested and initial EtOH intake (**Extended Figure 2-1**).

**Figure 2.**
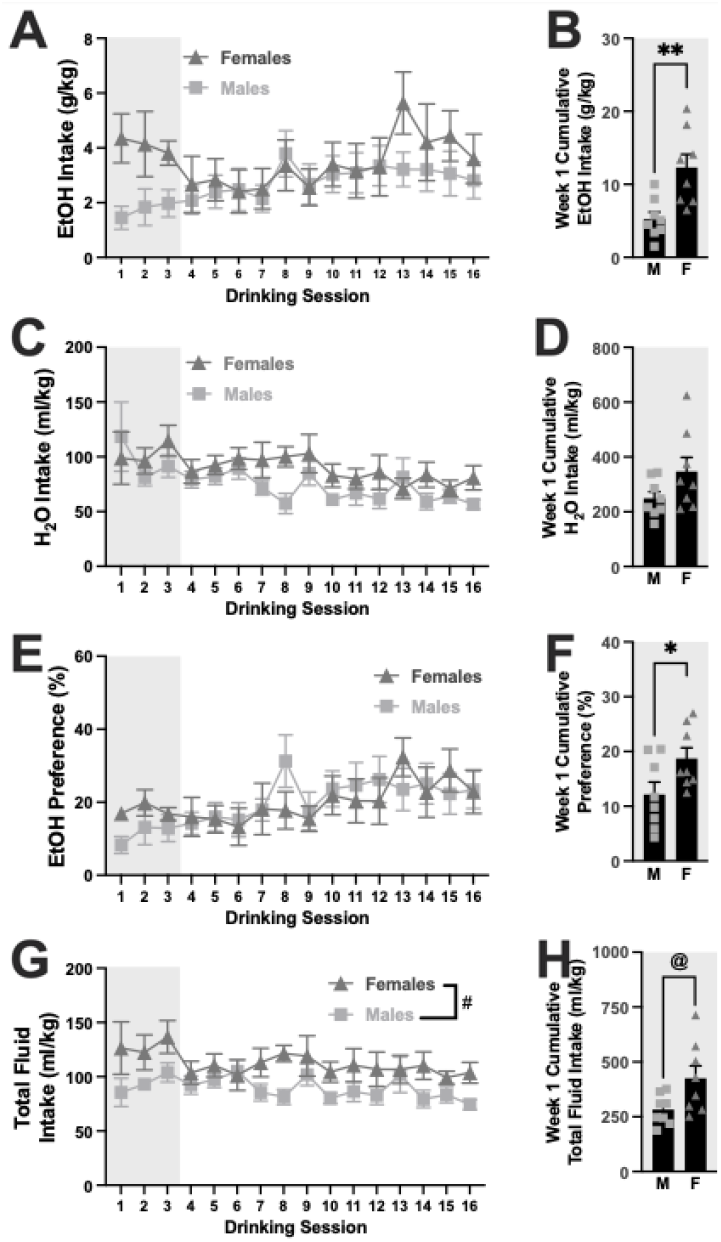
Voluntary EtOH drinking under an intermittent access paradigm. **A**) EtOH intake over time. **B**) Cumulative EtOH intake during the first week of EtOH access. **C**) Water intake over time. **D)** Cumulative water intake during the first week of EtOH access. **E**) EtOH preference over time. **F**) Cumulative EtOH preference during the first week of EtOH access. **G**) Total fluid intake over time. **H**) Cumulative total fluid intake during the first week of EtOH access. ** *p*<0.01, \**p*<0.01 (unpaired Student’s t test); & *p*<0.05 (2-way ANOVA, Main Effect of Sex); @ *p*<0.05 (unpaired Welch’s t test)

**Extended Figure 2-1.**
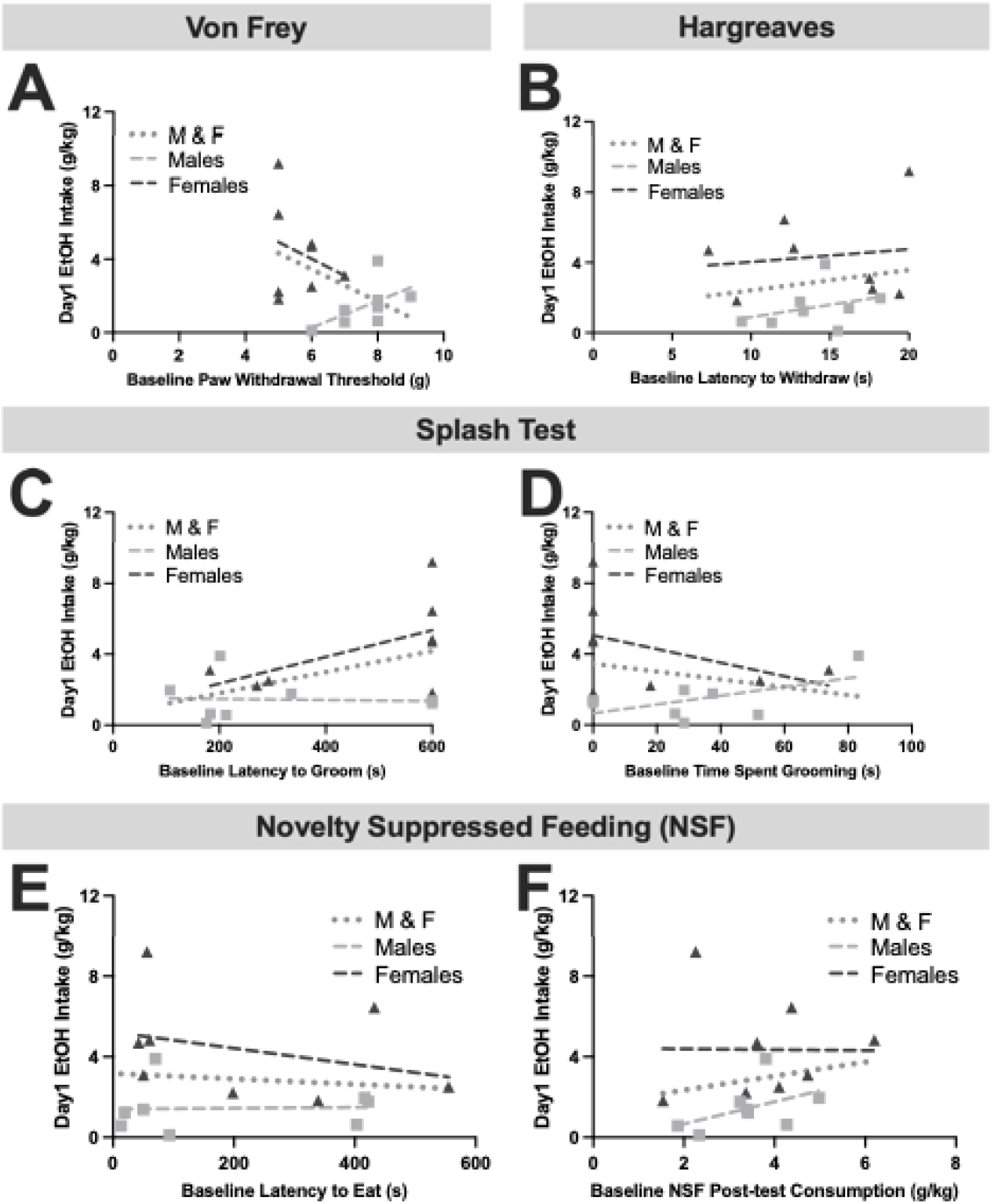
Behavioral measures did not correlate with first day EtOH intake in both males (M) and females (F). There was a trend where the rats with lower withdrawal thresholds drank more EtOH (**A**). Interestingly, there was no association between initial EtOH intake and basal thermal withdrawal (**B**), or basal latency to groom (**C**) and time spent grooming (**D**) in the splash test. There was also no significant relationship between initial EtOH intake and latency to eat (**E**) or post-test consumption (**F**) in the NSF test. <u>Pearson’s Correlations:</u> A) M&F: r(14)= −0.4742, *p*=0.0635; Males: r(6)=0.5673, *p*=0.1425; Females: r(6)=−0.2699, *p*=0.5179 B) M&F r(14)= 0.1836, *p*=0.4962; Males: r(6)=0.3344, *p*=0.4162; Females: r(6)=0.1388 *p*=0.7431 F) M&F: r(14)=0.1751, *p*=0.5167; Males: r(6)=0.4654, *p*=0.2451; Females: r(6)=−0.01082, *p*=0.9797 <u>Spearman’s Correlations:</u> C) M&F: r_s_(14)=0.3991, *p*=0.1266; Males: r_s_(6)=0.03593, *p*=0.9434; Females: r_s_(6)=0.4637, *p*=0.2619 D) M&F together: r_s_(14)=−0.2088, *p*=0.4355, Males: r_s_(6)=0.3234, *p*=0.4317; Females: r_s_(6)=−0.4364, *p*=0.3036 E) M&F: r_s_(14)=0.1177, *p*=0.6631; Males: r_s_(6)=0.3571; *p*=0.3894; Females: r_s_(6)=−0.2857, *p*=0.5008

### 3.4 EtOH Drinking & Associated Pain Sensitivity

After 3 weeks of EtOH access, the rats began post-drinking behavioral testing beginning with the Von Frey test of mechanosensitivity. There was a main effect of time, where paw withdrawal threshold decreased in all 4 groups^o^ (**Fig. 3A**), suggesting that all rats became more sensitive to mechanical stimuli over time. A main effect of EtOH^o^ also emerged, where EtOH-drinking rats had increased paw withdrawal thresholds, indicative of decreased mechanical sensitivity. Furthermore, there was an EtOH by time interaction, where Bonferonni posthoc analysis revealed that the decrease in paw withdrawal threshold over time was significantly stronger in water drinking rats compared to EtOH-drinking rats. Bonferroni posthoc analysis also indicated that the main effect of EtOH was driven predominantly by the post-test rather than pre-test, as there was no significant difference between water and EtOH-drinking rats at baseline. Finally, there was a main effect of sex^o^, where females exhibited decreased paw withdrawal thresholds suggesting increased mechanical sensitivity (consistent with **Fig. 1B**). Interestingly, paw withdrawal threshold did not correlate with previous day^p^ (**Fig. 3B**) or subsequent day EtOH intake^q^ (**Fig. 3C**).

**Figure 3.**
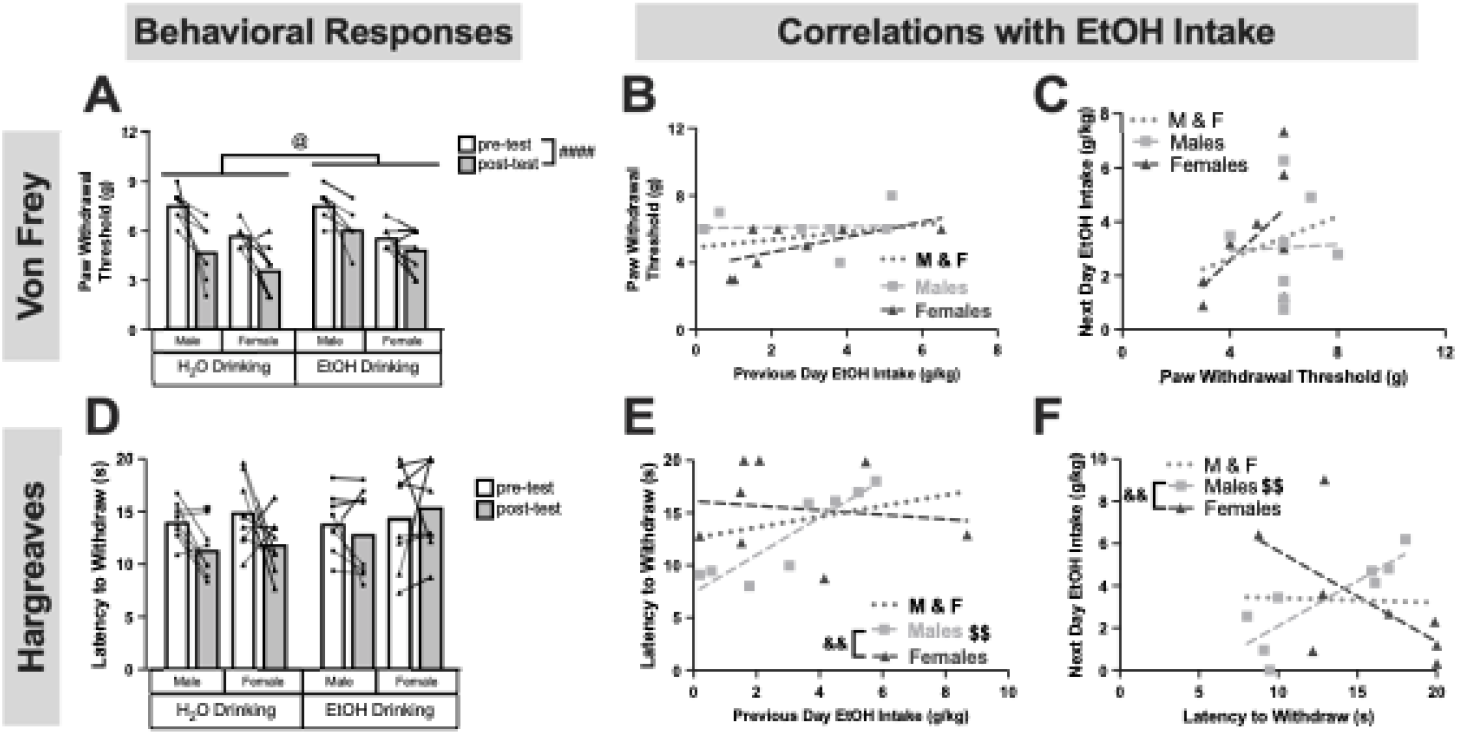
Mechanical and thermal sensitivity following chronic EtOH drinking in males and females. The Von Frey test was repeated and mechanical sensitivity (**A**) was correlated with previous day EtOH intake(**B**) and subsequent EtOH intake (**C**). The Hargreaves test was also repeated and thermal sensitivity (**D**) was correlated with previous day EtOH intake (**E**) and subsequent EtOH intake (**F**). #### *p*<0.0001 (3-way ANOVA, Main Effect of Time); @ *p*<0.05 (3-way ANOVA, Main effect of EtOH); $$*p*<0.01 (Pearson’s correlation); &&*p*<0.01 (Fisher’s r to z transformation). Pearson’s Correlations: **B)** M&F: r(14)=0.3155, *p*=0.2340; Males: r(6)=0.04155, *p*=0.9222; Females: r(6)=0.6262, *p*=0.0967 **C)** M&F: r(14)=0.2660, *p*=0.3193; Males: r(6)=0.03441, *p*=0.9355; Females: r(6)=0.5708, *p*=0.1395 **E)** M&F: r(14)=0.2846, *p*=0.2853; Males: r(6)=0.9043, *p*=0.0020; Females: r(6)=−0.1378, *p*=0.7448 **F)** M&F: r(14)=−0.03709, *p*=0.8915; Males: r(6)=0.8494, *p*=0.0075; Females: r(6)=−0.6227, *p*=0.0991

Next, the Hargreaves test was repeated. In this test, latency to withdraw the paw is used as an index of thermal sensitivity, where increased withdrawal latency is indicative of decreased thermal sensitivity. There were no significant effects of time, EtOH, or sex on withdrawal latency^r^ (**Fig. 3D**). Next, correlations examining the relationship between EtOH intake and withdrawal latency were performed. Previous day EtOH intake was directly correlated with withdrawal latency in males but not females^s^ (**Fig. 3E**). This association was also seen when examining next day EtOH intake, where withdrawal latency predicted subsequent EtOH intake in males but not females^t^ (**Fig. 3F**). The correlation of paw withdrawal threshold and EtOH intake in males as compared to females was statistically significant^s,t^ (**Fig. 3E-F**). Together, these findings reveal that increased EtOH intake is bidirectionally associated with decreased thermal sensitivity in a sex-specific manner. Finally, there were no changes in 24h EtOH intake produced by behavioral testing (**Extended Figure 3-1**).

**Extended Figure 3-1.**
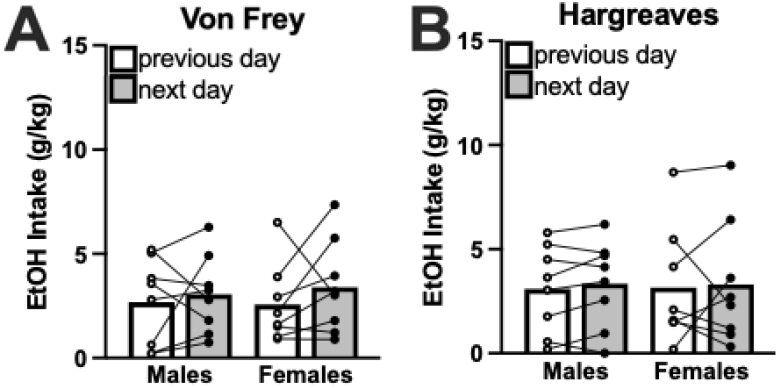
Pain testing does not affect EtOH Intake. There was no effect of Von Frey testing (**A**) or Hargreaves testing (**B**) on EtOH consumption. <u>2-way ANOVAs:</u> A) Sex × Day: F(1,14)=0.1799, *p*=0.6779; Sex: F(1,14)=0.01875, *p*=0.8930; Day: F(1,14)=1.244, *p*=0.2835 B) Sex × Day: F(1,14)=0.01586, *p*=0.9016; Sex: F(1,14)=5.719e005, *p*=0.9941; Day: F(1,14)=0.2869, *p*=0.6006

There was no effect of either EtOH drinking or sex on latency to groom^u^ (**Fig. 4A**) or total time spent grooming^v^ (**Fig. 4B**). However, a main effect of time emerged on total time spent grooming, where all rats spent significantly less time grooming on the posttest. This suggests that regardless of sex or history of EtOH drinking, rats exhibited increased depressive-like behavior over time.

**Figure 4.**
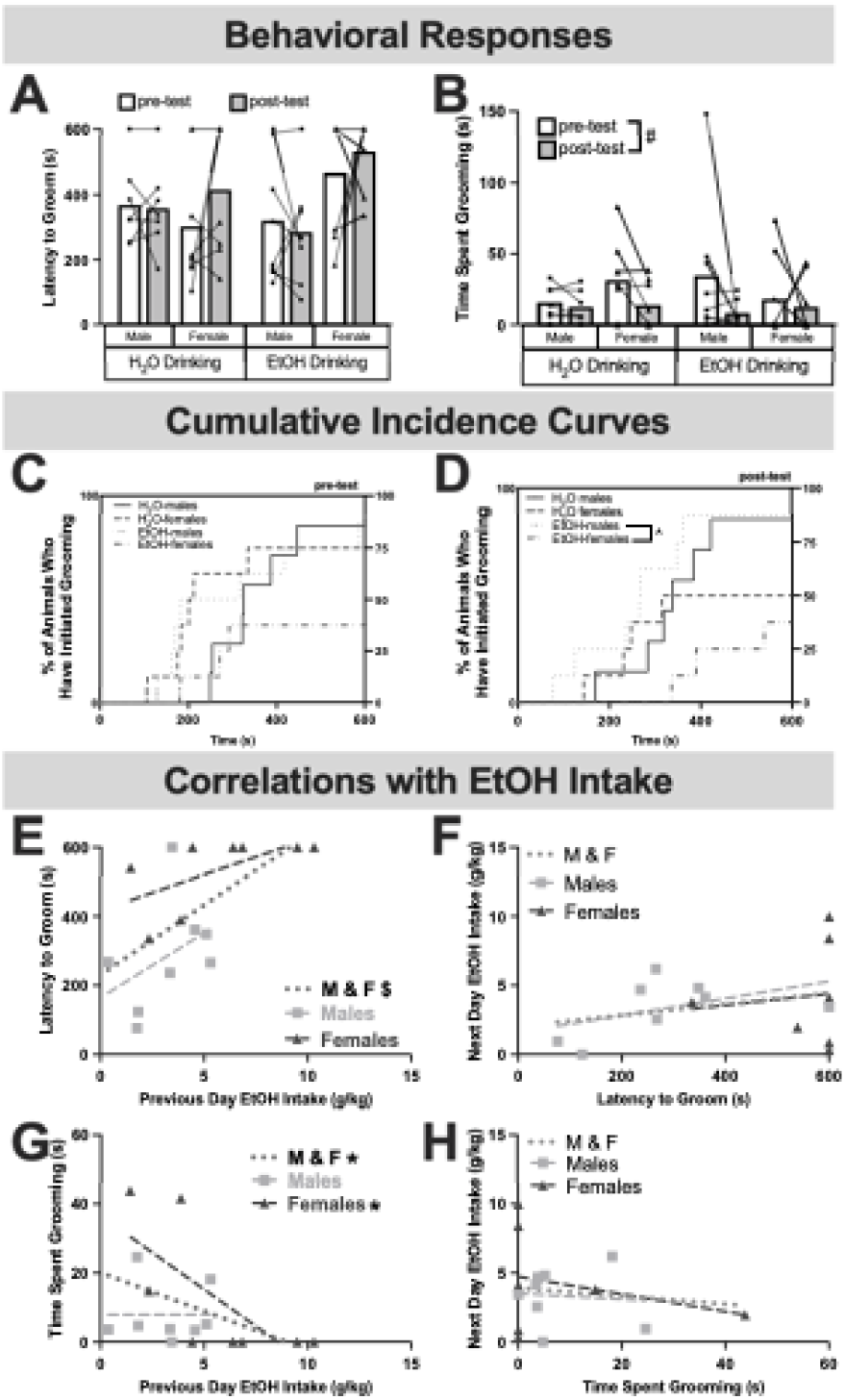
Anxiety- and depressive-like behavior in the splash test following chronic EtOH drinking in males and females. Latency to groom **(A)** and time spent grooming **(B)** were used as measured of anxiety- and depressive-like behavior. To account for animals that failed to groom within the 10min testing period, cumulative incidence curves for the pre-test and post-test are shown in **(C)**and **(D)**, respectively. The splash test was repeated and anxiety-like behavior was correlated with previous day EtOH intake (**E**) or subsequent EtOH intake (**F**). Depressive-like behavior in the splash test was correlated with previous day EtOH intake (**G**) or subsequent EtOH intake (**H**). # *p*<0.05 (3-way ANOVA, Main Effect of Time); ^*p*<0.05, Mantel-Cox log-rank test followed by Bonferroni post-hoc, $*p*<0.05 (Pearson’s correlation); *\* p*<0.05 (Spearman’s correlation) Pearson’s Correlations: **E)** M&F: r(14)=0.6108, p=0.0120; Males: r(6)=0.4120, *p*=0.3105; Females: r(6)=0.6172, *p*=0.1030 **F)** M&F: r(13)=0.2592, p=0.3508; Males: r(6)=0.4774, *p*=0.2269; Females: r(5)=0.1107, *p*=0.3132 Spearman’s Correlations: **G)** M&F: r_s_(14)=−0.5859, *p*=0.0193; Males: r_s_(6)=−0.8456, *p*=0.0179; Females: r_s_(6)=0.07143, *p*=0.0179 **H)** M&F: r_s_(13)=−0.02951, *p*=0.9180; Males: r_s_(6)=0.09524, *p*=0.8401; Females: r_s_(5)=−0.1782, *p*=0.7182

To account for animals that failed to groom during the 10 min test period, the cumulative incidence of grooming was plotted using the Kaplan-Meier survival curve followed by the Mantel-Cox log-rank test. There were no differences in cumulative incidence of grooming during the pre-test^w^ (**Fig. 4C**). However, there was a significant effect during the post-test^x^ (**Fig. 4D**), specifically where there was an increased cumulative incidence of grooming in EtOH-drinking males compared to EtOH-drinking females^x^. This suggests sex differences in anxiety-like behavior occur only in rats with a history of EtOH drinking (and not in EtOH-naïve rats).

To further investigate the association between EtOH intake and affective behavior, individual EtOH intake during the drinking session immediately before and after the Splash test were compared to latency to groom and total time spent grooming. Rats that consumed more EtOH prior to testing took significantly longer to begin grooming^y^ (**Fig. 4E**), an effect that was only seen when sexes were combined. This relationship was unidirectional, as latency to groom did not predict subsequent drinking^z^ (**Fig. 4F**). Total time spent grooming was also associated with EtOH intake prior to testing such that rats that consumed more EtOH groomed less^aa^ (**Fig. 4G**); however, this effect was primarily driven by females^w^. Additionally, there was no association between total time spent grooming and subsequent EtOH intake^ab^ (**Fig. 4H**) and the Splash test did not affect EtOH intake (**Extended Figure 4-1**). Together, these data suggest that previous day EtOH intake can predict both anxiety-like and depressive-like behavior in the Splash test.

**Extended Figure 4-1.**
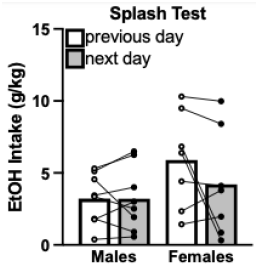
Splash test testing does not affect EtOH Intake. 2-way ANOVA: Sex × Day: *F*(1,13)=1.958, *p*=0.1852; Sex: *F*(1, 13)=1.883, *p*=0.1932; Day: F(1,13)=1.989, *p*=0.1819

The NSF test was also repeated to investigate changes in anxiety-like and motivated behavior following EtOH drinking. While there was no effect of sex or EtOH drinking on latency to eat^ac^, there was an effect of time^ac^ where all rats decreased their latency to eat on the second test (**Fig. 5A**). Post-test home-cage consumption was monitored for 10 minutes following the test and was used as a measure of motivation to eat. Compared to baseline, all the rats ate significantly less in their home-cage, suggesting a reduced motivation^ad^ (**Fig. 5B**). There were no differences in cumulative incidence of feeding initiation during baseline or after chronic drinking (**Extended Figure 5-1)** suggesting that while rats decreased their anxiety-like behavior in the NSF, there were no group differences.

**Figure 5.**
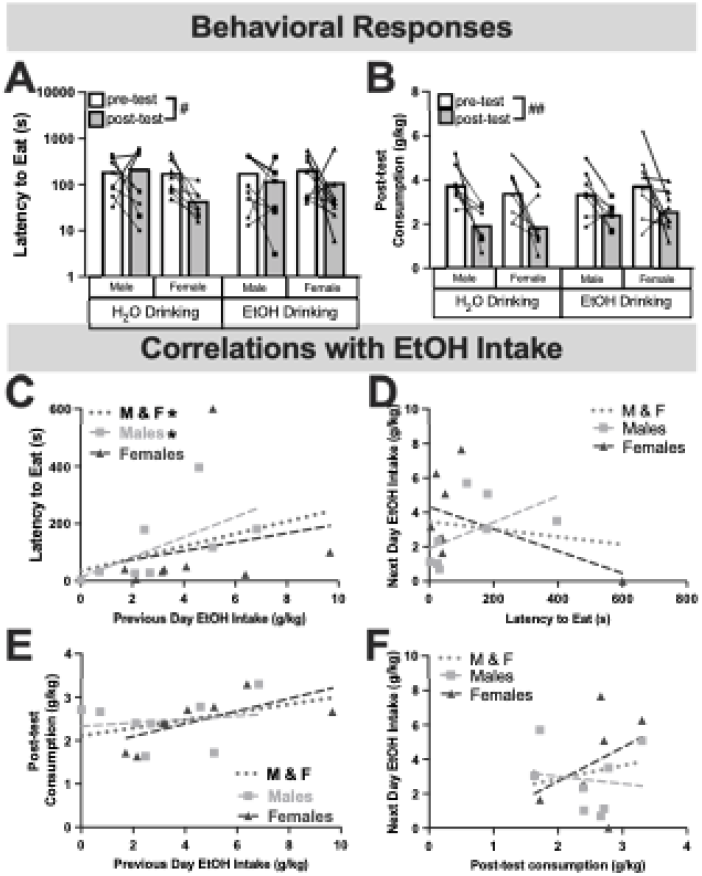
Anxiety-like behavior in the Novelty Suppressed Feeding test following chronic EtOH drinking in males and females. Latency to eat **(A)** and post-test consumption **(B)** were used as measured of anxiety-like and motivated behavior. Anxiety-like behavior was correlated with previous day EtOH intake (**C**) or subsequent EtOH intake (**D**). Motivated behavior was correlated with previous day EtOH intake (**F**) or subsequent EtOH intake (**G**). # *p*<0.05, ## *p*<0.01 (3-way ANOVA, Main Effect of Time); $*p*<0.05 (Pearson’s correlation); *\* p*<0.05 (Spearman’s correlation) Spearman’s Correlations: **C)** M&F: r_s_(14)=0.5647, *p*=0.0248; Males: r_s_(6)=0.7381, *p*=0.0458; Females: r_s_(6)=0.4286, *p*=0.2992 **D)** M&F: r_s_(14)=0.1912, *p*=0.4769; Males: r_s_(6)=0.6667, *p*=0.0831; Females: r_s_(6)=−0.1905, *p*=0.6646 Pearson’s Correlations: **E)** M&F: r(14)=0.4202, *p*=0.1051; Males: r(6)=0.1630, *p*=0.6997; Females: r(6)=−0.6833 *p*=0.0617 **F)** M&F: r(14)=0.1875, *p*=0.4867; Males: r(6)=−0.1263, *p*=0.7657; Females: r(6)=0.4296 *p*=0.2881

**Extended Figure 5-1.**
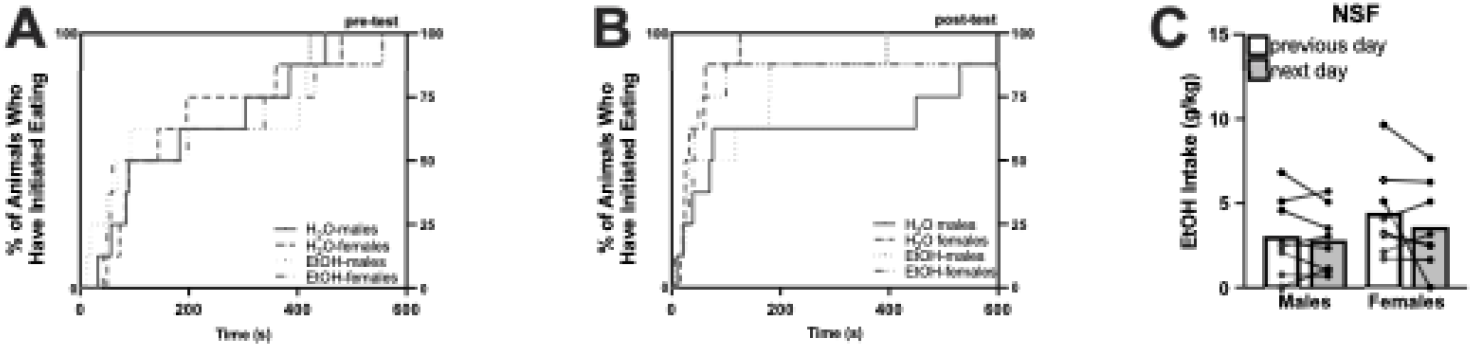
Cumulative occurrence of behavioral initiation in the NSF is not affected by EtOH drinking, nor is EtOH intake affected by NSF testing. There was no effect of group on cumulative occurrence of feeding initiation in the pre-test (**A**; Mantel-Cox log-rank test; X^2^=0.5295, df=3, *p*=0.9124) or post-test (**B**; Mantel-Cox log-rank test; X^2^=3.717, df=3, *p*=0.2937). Lastly, there was no effect of testing on EtOH intake (**C**; 2-way ANOVA; Sex × Day: *F*(1,13)=0.5462, *p*=0.4721; Sex: F(1,14)=1.901, *p*=0.3442; Day: F(1,14)=1.901, *p*=0.1896).

Associations between NSF behavioral measures and EtOH drinking were performed to examine potential individual effects of EtOH drinking on behavior. Rats that consumed more EtOH prior to the NSF test had increased latencies to eat,^ae^ and this effect was driven primarily by males (**Fig. 5C**). There were no significant correlations between latency to eat and subsequent EtOH intake^af^ (**Fig. 5D**). Post-test home-cage consumption did not correlate with previous day^ag^ (**Fig. 5E**) or subsequent EtOH intake^ah^ (**Fig. 5F**). Similar to the Splash test, there was no effect of testing on EtOH intake in either sex (**Extended Figure 5-1**). Together, these correlations suggest that EtOH intake is correlated with subsequent anxiety-like behavior on the NSF test.

### 3.7 EtOH drinking and Medial Prefrontal Cortex Activity

Following the NSF test, rats were allowed one final drinking session before brains were collected and the medial prefrontal cortex was stained for RFP (to identify CRF1+ cells) and cFos (a marker of neuronal activity). The medial prefrontal cortex was divided into prelimbic (PrL) and infralimbic (IL) cortices, based on the Paxinos and Watson atlas^54^.

When investigating PrL (**Fig. 6A**), there were no group differences in CRF1+ cells^ai^ (**Fig. 6B**), cFos+ cells^aj^ (**Fig. 6C**), total double-labeled cells^ak^ (**Fig. 6D**) or double-labeled cells as a percentage of total CRF1+ cells^al^ (**Fig 6E)**. Additionally, final session EtOH intake was not associated with any of these cell counts (**Extended Figure 6-1**). In the IL (**Fig. 6F**), there was no effect of sex or EtOH on CRF1+ cells^am^ (**Fig. 6G**). However, females had increased cFos+ cells^an^ (**Fig. 6H**). There were no significant differences in total double-labeled cells^ao^ but the sex difference seen in **Fig. 6H** persisted when double-labeled cells were expressed as %CRF1+ cells^ap^ (**Fig. 6J**). The number of CRF1+ cells was directly correlated with EtOH intake in females^aq^, but was inversely correlated with EtOH intake in males^aq^ (**Fig. 6K**). Additionally, the number of cFos+ cells was inversely correlated with EtOH intake in males only^ar^ (**Fig. 6L**). Consistent with the correlations seen thus far, total double-labeled cells was inversely correlated with EtOH intake in males only^as^ (**Fig. 6M**), however this correlation disappeared when double labeled cells were normalized to %CRF1+ cells^at^ (**Fig. 6N**). Taken together, these findings highlight differential effects of EtOH drinking on medial prefrontal cortex subdivisions, and suggest sex- and dose-dependent effects of EtOH on IL CRF1+ and cFos+ cells.

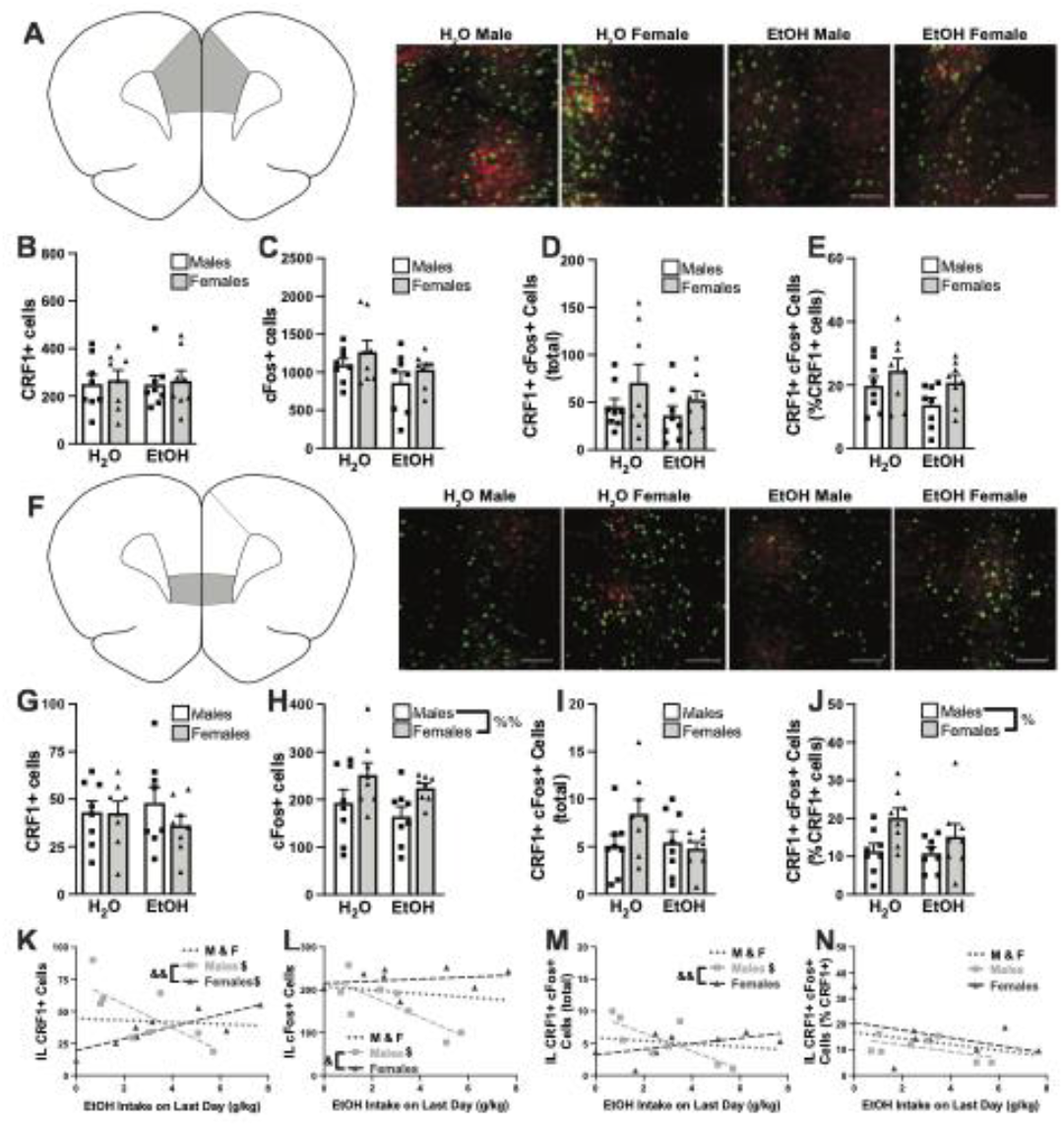
Prefrontal correlates of EtOH Drinking in males and females. Slices containing both the prelimbic cortex (PrL) and infralimbic cortex (IL) were stained for RFP (as a measure of CRF1+ neurons, shown in red) and cFos (as a measure of cellular activity, shown in green). PrL representative images **(A)** showing CRF1+ **(B)**, cFos+ **(C)** and total double labeled cells **(D)**. Double labeled cells were also expressed as % total CRF1+ cells (**E**). IL representative images **(F)**showing CRF1+ **(G)**, cFos+ **(H)** and total double labeled cells **(I)**. Double labeled cells were also expressed as % total CRF1+ cells (**J)**. We also correlated EtOH intake on the last day with IL labeling of CRF1+ cells **(K)**, cFos+ cells **(L)**,total double labeled cells **(M)** and double labeled cells as %CRF1+ **(N)**. Scale bars= 100μm; %%*p*<0.01 (2-way ANOVA; Main effect of Sex); $*p*<0.05 (Pearson’s correlation); &&*p*<0.01 (Fisher’s r to z transformation) Pearson’s Correlations: **K)** M&F: r(14)=−0.07998, *p*=0.7684; Males: r(6)=−0.7277, *p*=0.0407; Females: r(6)=0.8307, *p*=0.0106 **L)** M&F: r(14)=−0.1695, *p*=0.5403; Males: r(6)=−0.7775, *p*=0.0231; Females: r(6)=0.2192, *p*=0.6022 **M)** M&F: r(14)=−0.1909, *p*=0.4787; Males: r(6)=−0.7752, *p*=0.0238; Females: r(6)=0.5508, *p*=0.1571 **N)** M&F: r(14)=−0.3485, *p*=0.1858; Males: r(6)=−0.5444, *p*=0.1630; Females: r(6)=−0.4090, *p*=0.3144

**Extended Figure 6-1.**
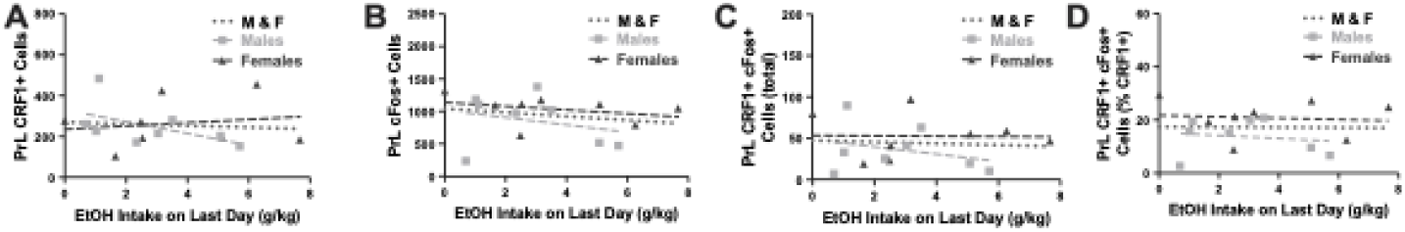
Correlations between EtOH intake and activity in the prelimbic cortex. EtOH intake did not correlate with CRF1+ expression (**A**), cFos+ cells (**B**), total double-labeled cells (**C**) or double-labeled cells when expressed as %CRF1+ cells (**D**). Spearman’s Correlations A) M&F: r_s_(14)=−0.1735, *p*=0.5193; Males: r_s_(6)=−0.5476, *p*=0.1710; Females: r_s_(6)=0.09524, *p*=0.8401 Pearson’s Correlations B) M&F: r(14)=−0.1997, *p*=0.4582; Males: r(6)=−0.25643, *p*=0.5271; Females: r(6)=−0.3325, *p*=0.4210), C) M&F: r(14)=−0.07430, *p*=0.7842; Males: r(6)=−0.2966, *p*=0.4757; Females: r(6)=−0.02222, *p*=0.9583 D) M&F: r(14)=−0.01427, *p*=0.9582; Males: r(6)=−0.1604, *p*=0.7043; Females: r(6)=−0.09422, *p*=0.82

## 4. DISCUSSION

The current study examined the dynamic relationship between affective states, pain sensitivity, and alcohol intake in male and female CRF1:cre: ^td^Tomato rats. While female rats displayed increased basal sensitivity to mechanical stimuli, there were no other sex differences in baseline behavior. Females consumed significantly more EtOH during the first week of intermittent access, but overall EtOH intake was not significantly different between males and females. EtOH drinking decreased mechanical sensitivity in both male and female rats, but mechanical sensitivity did not correlate with previous or subsequent EtOH intake. Conversely, there were no group effects of EtOH on thermal sensitivity; but in males, decreased thermal sensitivity was associated with increased EtOH intake before and after the Hargreaves test. There were no group effects of EtOH on anxiety- or depressive-like behavior; however, both anxiety- and depressive-like behavior were directly correlated with EtOH intake prior to testing, but were not associated with subsequent EtOH intake. Additionally, all groups displayed increased mechanical sensitivity and depressive-like behavior as well as decreased motivated behavior compared to baseline levels. Finally, we investigated potential effects of EtOH intake on the activity of CRF1-containing neurons in the prelimbic and infralimbic prefrontal cortex. While there were no significant group effects of alcohol on overall or CRF1-specific activity, there were significant sex-specific associations between EtOH intake and activity in the infralimbic region. These findings provide new insight into individual differences in EtOH intake and its association with affective state, pain sensitivity, and neuronal activation of CRF1-containing neurons in key regions of the prefrontal cortex.

One important goal of this study was to examine affective behavior and EtOH drinking in male and female CRF1:cre: ^td^Tomato rats. There was no effect of sex on basal affective behavior with the exception of mechanical sensitivity, where female rats displayed increase mechanical sensitivity. One previous study examined pain sensitivity in male and female CRF1:cre: ^td^Tomato rats; however, data were not reported by sex^52^. Studies involving Wistar rats (the background strain for the CRF1:cre:^td^Tomato rats) also found no sex differences in baseline anxiety- and depressive-like behavior in the Splash Test^56, 57^. Sex differences in the NSF test in Wistar rats are less clear: one study found females exhibited increased anxiety-like behavior^58^; however, other studies report no sex differences^59–61^. Following baseline behavioral testing, rats were allowed to voluntarily drink EtOH under an intermittent access paradigm. In this study, male and female CRF1:cre: ^td^Tomato rats drank 2-4g/kg EtOH with an approximate 15-30% EtOH preference, consistent with studies in Wistar rats^62–68^. While there were no sex differences in EtOH intake throughout the experiment, females drank significantly more EtOH during the first week. Studies comparing male and female EtOH intake under intermittent access conditions in Wistar rats are limited and inconclusive: some researchers found females drank more EtOH whereas others found no sex differences^67, 69^. Future studies will examine EtOH drinking in other contexts, such as binge-drinking or drinking following abstinence from EtOH. The studies presented here expand our knowledge of baseline behavior and EtOH intake in male and female CRF1:cre: ^td^Tomato rats.

Although the behavioral tests used in these studies examine different aspects of affective behavior, one common theme emerged in the post-drinking tests: negative affect increased in both EtOH drinking animals as well as EtOH-naïve animals. Specifically, we found increased negative affect in both sexes in the Von Frey test (decreased withdrawal thresholds, suggesting increased mechanical sensitivity), Splash test (decreased time spent grooming, suggestive of decreased motivation), and NSF test (decreased post-test consumption, suggestive of decreased motivated behavior). Our study aimed to investigate the directionality of the relationship between negative affect and EtOH drinking, and therefore having baseline measures were crucial. However, affective tests are rarely repeated in the same subjects; thus, there are limited findings regarding the effects of time and re-testing on behavior. In one study, researchers found that male rats, but not female rats, spent significantly less time grooming on the second splash test^70^. While the current findings indicate decreased motivation regardless of sex, strain differences may account for this discrepancy. One potential explanation for the increased negative affect is increased stress due to prolonged social isolation. Indeed, prolonged social isolation during adulthood increases pain and negative affect in both humans and animals^71–78^. Together, these studies add to the literature by suggesting prolonged social isolation increases mechanical sensitivity and depressive-like behavior in both males and females regardless of EtOH exposure.

Many studies have reported increased negative affect following chronic EtOH drinkingt^36, 79–82^. However, there were no group effects of EtOH on thermal sensitivity or anxiety- and depressive-like behavior in the studies presented here. One potential explanation is that rats in the present study only consumed EtOH for 3-4 weeks before behavioral testing and thus did not consume enough EtOH to produce behavioral deficits. Studies in Wistar rats report escalated intake after ~20 EtOH drinking sessions^64, 83, 84^; rats in this study only drank for 16 sessions. Others have found that various aspects of EtOH withdrawal (including increased anxiety-like behavior, seizures, and EtOH intake during withdrawal) increase with repeated EtOH withdrawal cycles^85–90^. Another possibility is that the environment was too anxiogenic, creating a ceiling effect on anxiety-like behavior. Recent studies have developed methods to test anxiety-like behavior in the home-cage^91^ to remove this potential confound and will be considered in the future.

Despite the lack of group effects, there were significant individual correlations between affective behavior and EtOH intake, thus highlighting the importance of examining individual differences in the context of EtOH-related behaviors. More specifically, this study found previous EtOH correlated with thermal sensitivity as well as anxiety- and depressive-like behavior. There was an inverse relationship between EtOH intake and thermal sensitivity, where males that consumed more EtOH were less sensitive to thermal stimuli and males with increased thermal sensitivity consumed less EtOH on the next day. These findings are in accordance with a previous study reporting that EtOH withdrawal increases nociceptive threshold^92^. However, it is important to note that other studies have demonstrated EtOH-induced decreases in pain threshold ^93–97^. One potential explanation is that EtOH-induced peripheral neuropathy blunted the ability of the rats to feel the thermal stimulus. This would be consistent with the group effect of EtOH on mechanical sensitivity, where the EtOH group had increased paw withdrawal thresholds, corresponding to decreased mechanical sensitivity. An alternative possibility is that CRF1:cre: ^td^Tomato rats may be resistant to the analgesic properties of EtOH; however, additional studies are required to examine this in more detail. In addition to associations with thermal sensitivity, previous day EtOH intake was also associated with anxiety- and depressive-like behavior in the Splash test and NSF test. These findings are consistent with previous studies that demonstrate increased fear-potentiated startle in various genetic lines of alcohol preferring rats^98^ and mice^99^. While all of these correlations held true when both sexes were analyzed together, analyzing the sexes separately revealed that the EtOH intake associations with anxiety-like behavior in the NSF were primarily driven by males whereas EtOH intake associations with depressive-like in the Splash test were primarily driven by females. These findings suggest test-specific contributions of sex in EtOH-induced affective behavior. Interestingly, anxiety-like and depressive-like behavior were unable to predict subsequent EtOH intake. Other studies have mixed findings regarding associations of anxiety-like behavior and subsequent drinking with significant correlations found in some studies^9–11^ but not others^17, 100–102^. This work expands on these findings by demonstrating that neither baseline nor post-drinking anxiety- or depressive-like behavior predicts EtOH intake in male or female rats.

The effects of alcohol on affective behaviors are known to be mediated by key brain regions including the medial prefrontal cortex. Many studies fail to differentiate the prelimbic (PL) and infralimbic (IL) subdivisions of this region, but it is clear that these regions often play opposing roles in mediating behavior. For example, dopamine blockade in the PL reduces place conditioning without affecting cue conditioning but dopamine blockade in the IL reduces cue conditioning without affecting place conditioning^103^. While both PL and IL are active during cue-induced reinstatement of EtOH seeking, only ablation of activity-dependent IL neurons inhibited EtOH seeking^104^. Another study found EtOH drinking during adolescence increased Arc immunoreactivity in PL (but not in IL) whereas EtOH drinking during adulthood increased Arc immunoreactivity in IL (but not PL)^105^. While the lack of effect in PrL is consistent with the present findings, the effects in IL were divergent in that our study found decreased activity in the IL whereas their study found increased IL activity. In addition to species differences (mice vs rats), the timing of tissue collection may explain the differences seen. In the referenced study, brains were collected 25d after the last drinking session, but in the studies presented here, brains were collected 24h after the last drinking session. It is possible that increased length of forced abstinence results in compensatory upregulation of neuronal activity. Indeed, many studies have demonstrated opposing effects during short-term abstinence (~24h) and long-term abstinence (>7d)^106, 107^.PL and IL also exert opposing effects on stress reactivity. For example, PL lesions result in augmented stress responses whereas IL lesions result in attenuated stress responses^108^. These are just a few studies that highlight the necessity to investigate PL and IL as separate regions, rather than combining them into one. In the studies presented here, there were no group differences or individual differences in the effects of chronic EtOH drinking on neuronal activation (cFos+ cells) or activation of

CRF1+ cells (double labeled cells) in the PL. However, individual differences emerged when examining the association between EtOH intake and cFos+ cells as well as EtOH intake and activated CRF1+ cells in the IL. Not only were these associations region-specific, but also sex-dependent as the correlations only occurred in male rats. There are limited studies investigating the effects of EtOH drinking in CRF1+ cells in the PL and IL subdivisions of the PFC. An important consideration is that most prior studies examined changes in CRF1 transcript levels; however, our studies examined the effects of EtOH on cells that contain CRF1 and can thus not be compared directly. One important study found that EtOH vapor withdrawal decreases excitability and excitatory neurotransmission in CRF1+ neurons in the prefrontal cortex^109^; however, this study was only performed in male mice and did not separate PL from IL. Collectively, these studies highlight the relevance of including both sexes and sub-regions while examining the effects of chronic EtOH drinking.

Together, our results illustrate the complex interplay between affective state, EtOH drinking, and the role of prefrontal cortex CRF1+ neurons in mediating these behaviors. Moreover, these data highlight the importance of examining individual differences in addition to group averages when investigating AUD and affective states. There are currently only 4 FDA-approved medications to treat AUD, and persistent issues with efficacy and compliance complicate effective treatment. Clinical trials have demonstrated differential efficacies of medications when stratified by factors such as sex, age of AUD onset, severity of AUD, and impulsivity levels^110, 111^. However, comorbidity of AUD and affective disorders remains an underexamined issue. Thus, additional studies that consider individual differences in affective behavior and population-specific neurobiological changes in the context of AUD are warranted.

## 13. Acknowledgements

We thank Jasmine J Jahad for their help in early intermittent access drinking experiments.

