## Supplemental Materials for "Chronic Alcohol Drinking Drives Sex-Specific Differences in Affective Behavior and Medial Prefrontal Cortex Activity in CRF1:Cre:Tdtomato Transgenic Rats"

**Effects of Chronic Alcohol Consumption on Affective Behavior and Medial Prefrontal Cortex Activity in Male and Female CRF1:cre:tdTomato Transgenic Rats**

Quadir SG^1^, Arleth GM^1^, Cone MG^1^, High MW^1^, Ramage MC^1^, Effinger DP^2^, Echeveste-Sanchez M^1^, Herman MA^1,2^

**Supplementary Figures**

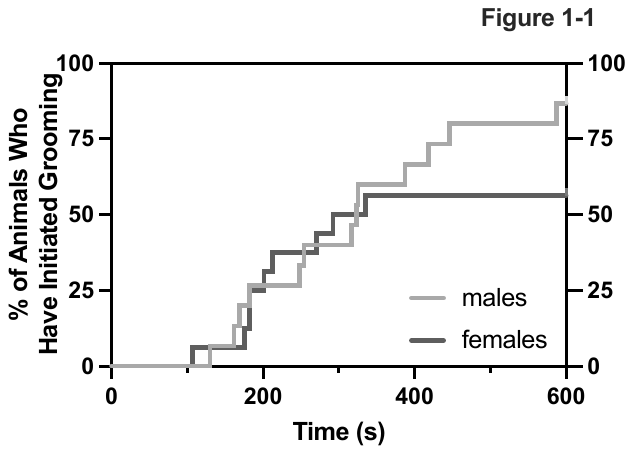

### Extended Figure 1-1

**Cumulative Incidence of Grooming Initiation in the Splash Test.** Even when accounting for the rats that never groomed (rather than artificially assigning them a maximum value or excluding them from studies), there was no sex difference in grooming initiation (Mantel-Cox log-rank test: χ²=1.430, df=1, *p*=0.2318). There was also no sex difference in total time spent grooming (**C;** Mann-Whitney test, *p*=0.8901).

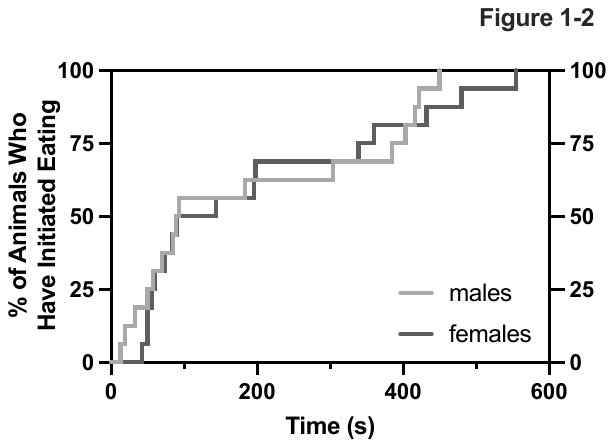

### Extended Figure 1-2

**Cumulative Incidence of Feeding Initiation in the Novelty Suppressed Feeding Test.** There was no effect of sex on cumulative occurrence of feeding initiation (**B**; Mantel-Cox log-rank test, χ²=0.3366, df=1, *p*=0.5618).

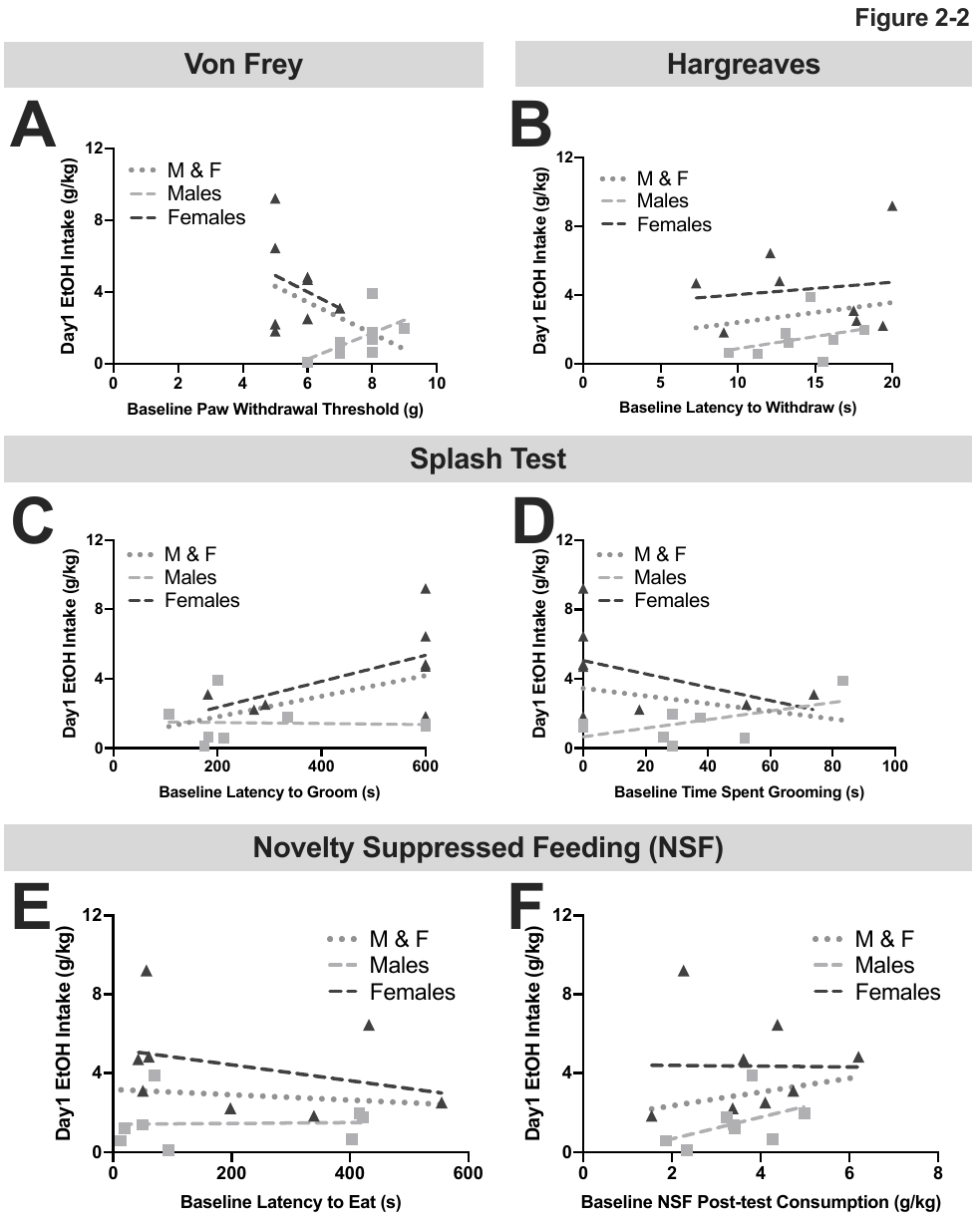

### Extended Figure 2-1

**Behavioral measures did not correlate with first day EtOH intake in both males (M) and females (F).** There was a trend where the rats with lower withdrawal thresholds drank more EtOH (**A**). Interestingly, there was no association between initial EtOH intake and basal thermal withdrawal (**B**), or basal latency to groom (**C**) and time spent grooming (**D**) in the splash test. There was also no significant relationship between initial EtOH intake and latency to eat (**E**) or post-test consumption (**F**) in the NSF test.

Pearson’s Correlations:

A) M&F: r(14)= -0.4742, *p*=0.0635; Males: r(6)=0.5673, *p*=0.1425; Females: r(6)=-0.2699, *p*=0.5179

B) M&F r(14)= 0.1836, *p*=0.4962; Males: r(6)=0.3344, *p*=0.4162; Females: r(6)=0.1388 *p*=0.7431

F) M&F: r(14)=0.1751, *p*=0.5167; Males: r(6)=0.4654, *p*=0.2451; Females: r(6)=-0.01082, *p*=0.9797

Spearman’s Correlations:

C) M&F: r_s_(14)=0.3991, *p*=0.1266; Males: r_s_(6)=0.03593, *p*=0.9434; Females: r_s_(6)=0.4637, *p*=0.2619

D) M&F together: r_s_(14)=-0.2088, *p*=0.4355, Males: r_s_(6)=0.3234, *p*=0.4317; Females: r_s_(6)=-0.4364, *p*=0.3036

E) M&F: r_s_(14)=0.1177, *p*=0.6631; Males: r_s_(6)=0.3571; *p*=0.3894; Females: r_s_(6)=-0.2857, *p*=0.5008

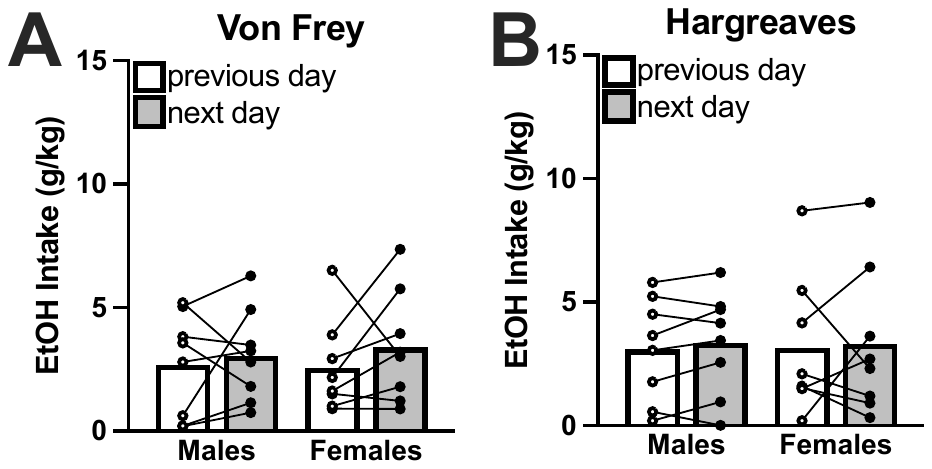

### Extended Figure 3-1

**Pain testing does not affect EtOH Intake.** There was no effect of Von Frey testing (**A**) or Hargreaves testing (**B**) on EtOH consumption.

2-way ANOVAs:

A) Sex × Day: F(1,14)=0.1799, *p*=0.6779; Sex: F(1,14)=0.01875, *p*=0.8930; Day: F(1,14)=1.244, *p*=0.2835

B) Sex × Day: F(1,14)=0.01586, *p*=0.9016; Sex: F(1,14)=5.719e005, *p*=0.9941; Day: F(1,14)=0.2869, *p*=0.6006

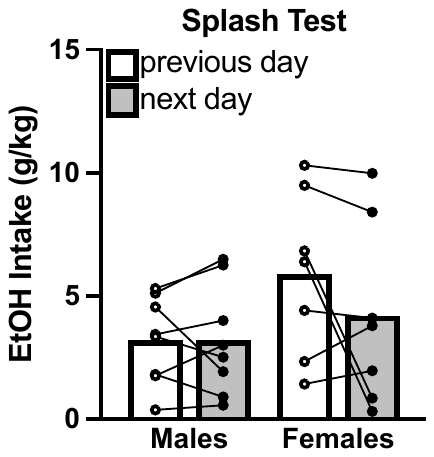

### Extended Figure 4-1

**Splash test testing does not affect EtOH Intake.** 2-way ANOVA: Sex × Day: *F*(1,13)=1.958, *p*=0.1852; Sex: *F*(1,13)=1.883, *p*=0.1932; Day: F(1,13)=1.989, *p*=0.1819

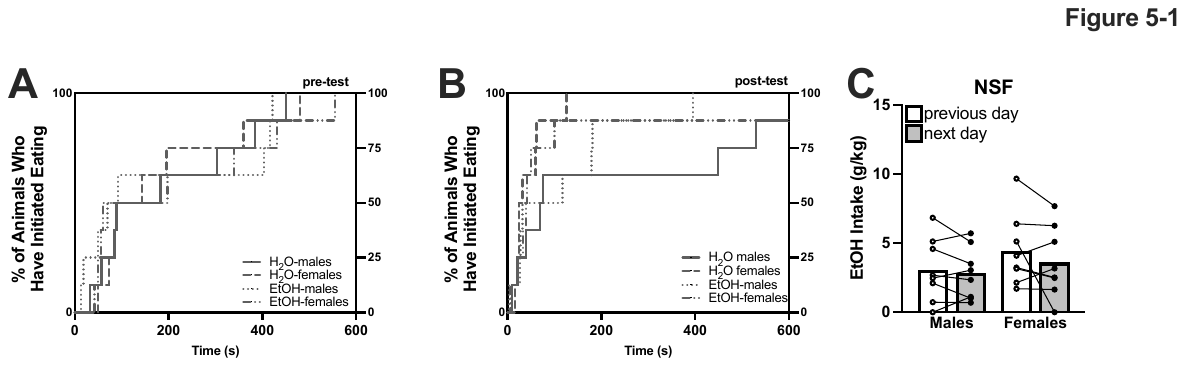

### Extended Figure 5-1

**Cumulative occurrence of behavioral initiation in the NSF is not affected by EtOH drinking, nor is EtOH intake affected by NSF testing.** There was no effect of group on cumulative occurrence of feeding initiation in the pre-test (**A**; Mantel-Cox log-rank test; χ²=0.5295, df=3, *p*=0.9124) or post-test (**B**; Mantel-Cox log-rank test; χ²=3.717, df=3, *p*=0.2937). Lastly, there was no effect of testing on EtOH intake (**C**; 2-way ANOVA; Sex × Day: *F*(1,13)=0.5462, *p*=0.4721; Sex: F(1,14)=1.901, *p*=0.3442; Day: F(1,14)=1.901, *p*=0.1896).

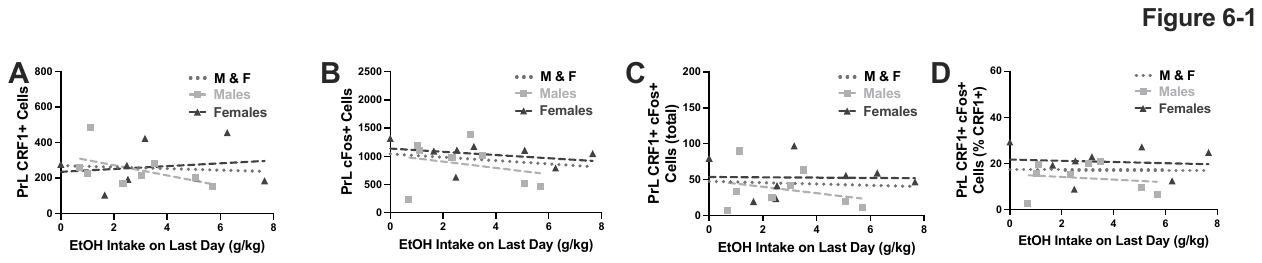

### Extended Figure 6-1

**Correlations between EtOH intake and activity in the prelimbic cortex.** EtOH intake did not correlate with CRF1+ expression (**A**), cFos+ cells (**B**), total double-labeled cells (**C**) or double-labeled cells when expressed as %CRF1+ cells (**D**)**.**

Spearman’s Correlations

A) M&F: r_s_(14)=-0.1735, *p*=0.5193; Males: r_s_(6)=-0.5476, *p*=0.1710; Females: r_s_(6)=0.09524, *p*=0.8401

Pearson’s Correlations

B) M&F: r(14)=-0.1997, *p*=0.4582; Males: r(6)=-0.25643, *p*=0.5271; Females: r(6)=-0.3325, *p*=0.4210),

C) M&F: r(14)=-0.07430, *p*=0.7842; Males: r(6)=-0.2966, *p*=0.4757; Females: r(6)=-0.02222, *p*=0.9583
D) M&F: r(14)=-0.01427, *p*=0.9582; Males: r(6)=-0.1604, *p*=0.7043; Females: r(6)=-0.09422, *p*=0.82
